# Group B streptococcal membrane vesicles induce proinflammatory cytokine production and are sensed in an NLRP3 inflammasome-dependent mechanism in human macrophages

**DOI:** 10.1101/2022.08.10.503555

**Authors:** Cole R. McCutcheon, Jennifer A. Gaddy, David M. Aronoff, Shannon D. Manning, Margaret G. Petroff

**Affiliations:** Department of Microbiology and Molecular Genetics, Michigan State University, East Lansing, MI, 48823; Department of Medicine, Division of Infectious Disease, Vanderbilt University Medical Center, Nashville, TN, 37232; Department of Pathology, Microbiology, and Immunology, Vanderbilt University Medical Center, Nashville, TN, 37232; Tennessee Valley Healthcare System, Department of Veterans Affairs, Nashville, TN; Department of Medicine, Indiana University School of Medicine, Indianapolis, IN, 46202; Department of Pathobiology and Diagnostic Investigation, Michigan State University, East Lansing, MI, 48823

## Abstract

Group B *Streptococcus* (GBS) is a major cause of fetal and neonatal mortality worldwide. Many of the adverse effects associated with invasive GBS are associated with inflammation that leads to chorioamnionitis, preterm birth, sepsis, and meningitis; therefore, understanding bacterial factors that promote inflammation is of critical importance. Membrane vesicles (MVs), which are produced by many pathogenic and non-pathogenic bacteria, may modulate host inflammatory responses. In mice, GBS MVs injected intra-amniotically can induce preterm birth and fetal death. Although it is known that GBS MVs induce large-scale leukocyte recruitment into infected tissues, the immune effectors driving these responses are unclear. Here, we hypothesized that macrophages respond to GBS-derived MVs by producing proinflammatory cytokines and are recognized through one or more pattern recognition receptors. We show that THP-1 macrophage-like cells produce high levels of neutrophil- and monocyte-specific chemokines in response to MVs derived from different clinical isolates of GBS. Interleukin (IL)-1β was significantly upregulated in response to MVs, which was independent of NF-kB signaling but dependent on both caspase-1 and NLRP3. These data indicate that MVs contain one or more pathogen-associated molecular patterns that can be sensed by the immune system. Furthermore, this study identifies the NLRP3 inflammasome as a novel sensor of GBS MVs. Our data additionally indicate that MVs may serve as immune effectors that can be targeted for immunotherapeutics, particularly given that similar responses were observed across this subset of GBS isolates.

## INTRODUCTION

Group B *Streptococcus* (GBS) is an opportunistic pathogen that colonizes the vaginal or rectal tract of ~30% of women (1). While maternal colonization is often asymptomatic, GBS can cause severe infections in pregnant women and neonates (1). Pregnancy- and neonatal-associated GBS infections are often characterized by pathologies exhibiting a high degree of inflammation. During pregnancy, this can present as placental villitis and preterm birth, whereas in neonates, GBS can cause meningitis and sepsis (2–4). Despite the high colonization frequencies in mothers, only a fraction of women and their neonates develop these threatening infections. The reasons for this discrepancy, however, are incompletely characterized.

We and others have postulated that strain variation contributes to the discrepancy in disease outcome. Indeed, specific phylogenetic lineages of GBS, which are defined by multilocus sequence typing (MLST) are more likely to cause neonatal infections (5–7). Notably, sequence type (ST)-17 strains are more commonly associated with invasive neonatal infections (5, 8, 9), whereas ST-1 strains are associated with invasive disease in adults (10). Conversely, ST-12 strains have been linked to asymptomatic maternal colonization (11). We demonstrated that ST-17 strains elicit stronger proinflammatory immune responses and persist longer inside macrophages than other strains (12, 13). Interestingly, we also found that ST-1 and ST-17 strains induce stronger activation of the proinflammatory transcription factor NF-kB compared to ST-12 strains (13). While ST-17 strains were previously found to have unique virulence gene profiles relative to other lineages, the specific bacterial factor(s) promoting these altered inflammatory responses are not fully understood (14–16).

Recently it was reported that GBS produces membrane vesicles (MVs) that can induce substantial recruitment of neutrophils and lymphocytes into murine extraplacental membranes, which mimicked GBS-associated chorioamnionitis in humans (17, 18). In support of this finding, GBS MVs were shown to induce production of the neutrophil chemokine CXCL1 in a murine model of *in utero* infection (17, 19), which has been shown in other GBS infection models (20, 21). Further, we recently reported that GBS MV production varies in abundance and protein composition across STs (17, 19, 22). More specifically, several immunomodulatory virulence factors, including hyaluronidase, C5a peptidase, and sialidase were highly and differentially abundant across STs (22). Together these data indicate that MVs promote proinflammatory immune responses; however, no prior studies have comprehensively examined the mechanisms by which human leukocytes respond to GBS MVs.

As sentinel leukocytes at the maternal fetal interface, macrophages play an important role in shaping immune responses. At the maternal-fetal interface macrophages make up 20-30% of leukocytes (23) and play pivotal roles in fertility (24), placental function (25), and host-pathogen interactions at the maternal-fetal interface (26–28). The THP-1 monocytic leukemia cell line can be differentiated with phorbol esters into macrophage-liked cells (29) and serve as a model system to evaluate host responses to GBS (12, 30). Using this model, we previously showed that THP-1 cells produce high levels of proinflammatory cytokines in response to GBS. Interestingly, several cytokines displayed lineage-specific inflammatory responses, with ST-17 strains eliciting a more potent inflammatory response compared to other lineages (13). Here, we examined macrophage responses to GBS MVs isolated from a diverse set of strains and found that these MVs induce the production of proinflammatory cytokines and chemokines. We also identified NLRP3 as a sensor of GBS derived MVs. In all, this study has expanded our current understanding of how host cells respond to GBS MVs. Additionally, by identifying the pathways upregulated by MVs, we have identified the proinflammatory pathways and receptors that could be used as potential immunotherapeutic targets.

## METHODS

### Bacterial Strains and Culture

GBS strains GB0037 (GB37), GB0411 (GB411), GB0653 (GB653), and GB1455 were isolated as described previously (31, 32). The invasive isolates GB37, GB411, and GB1455, were isolated from the blood or cerebrospinal fluid of infants with early onset GBS disease (31), while the colonizing strain GB653 was isolated from vaginal/rectal swabs collected from an asymptomatically colonized mother before childbirth (32). These isolates were previously characterized by MLST and capsular serotyping (9, 11). The GBS strains analyzed here represent colonizing and invasive isolates belonging to each of three common STs: ST-1 (GB37), ST-12 (GB1455 and GB653), and ST-17 (GB411). Strains were cultured using Todd-Hewitt Broth (THB) or Todd-Hewitt Agar (THA) (BD Diagnostics, Franklin Lakes, New Jersey, USA) overnight at 37°C with 5% CO_2_.

### Membrane vesicle isolation

MVs were isolated as previously described (22). Briefly, overnight THB cultures were diluted 1:50 into fresh broth and grown to late logarithmic phase (optical density (OD)_600_ = 0.9). Cultures were centrifuged at 2000 x g for 20 minutes at 4°C. Supernatants were collected and re-centrifuged at 8500 x g for 15 minutes at 4°C, followed by filtration through a 0.22μm filter and concentration using Amicon Ultra-15 centrifugal filters (10 kDa cutoff) (MilliporeSigma, Burlington, MA, USA). Concentrated supernatants were subjected to ultracentrifugation for 2 hours at 150,000 x g at 4°C. Pellets were resuspended in PBS and purified using qEV Single size exclusion columns (IZON Science, Christchurch, New Zealand) per the manufacturer’s instructions. MV fractions were collected and re-concentrated using the Amicon Ultra-4 centrifugal filters (10 kDa cutoff) (MilliporeSigma, Burlington, Massachusetts, USA) and brought to a final volume of 100 μL in PBS. MVs were aliquoted and stored at −80°C until further use.

### Nanoparticle Tracking Analysis

MVs were quantified via nanoparticle tracking analysis using a NanoSight NS300 (Malvern Panalytical Westborough, MA, USA) equipped with an automated syringe sampler as described previously (22, 33, 34). For each sample, MVs were diluted in PBS (1:100 – 1:1000) and injected with a flow rate of 50. Once loaded, five 20-second videos were recorded at a screen gain of 1 and camera level of 13, which were analyzed at a screen gain of 10 and a detection threshold of 4 after capture. Data were subsequently exported to a CSV file for analysis using the R package tidyNano (33).

### THP-1 Cell Culture

THP-1 cells (TIB-202) were obtained through ATCC (Manassas, VA) and stored according to vendor guidelines (35). Briefly, cells were cultured in RPMI 1640 (Gibco, ThermoFisher, Waltham, MA) supplemented with L-Glutamine, 10% fetal bovine serum (FBS), and 1% antibiotic-antimycotic (100 μg/mL Streptomycin, 0.25 ug/mL Amphotericin B, & 100 U/mL Penicillin; Gibco, ThermoFisher, Waltham, MA) as previously described (12, 13). For experiments, THP-1 cells were only utilized until passage 10. When indicated, THP-1 monocytes were differentiated into macrophages using phorbol 12-myristate 13-acetate (PMA) as previously described (12, 13). Cells were differentiated in RPMI (without phenol red) supplemented with L-Glutamine, 2% FBS and 100 nM PMA for 24 hours prior to experimentation (12, 13).

For experiments using GBS treated cells, THP-1 cells were washed twice with PBS prior to infection. The bacteria were resuspended in RPMI and added to the THP-1 cells at a multiplicity of infection (MOI) of 10 bacteria per cell. Cells were incubated for 1 hour and the media was subsequently aspirated. Cells were washed thrice with PBS and fresh RPMI with L-Glutamine (no phenol red) containing 2% FBS, 100 nM PMA, penicillin (5 μg/mL) and gentamicin (100μg/mL) was added (termed RPMI 2/0). Cells were incubated for an additional 24 hours. For MV treatment, cells were washed twice, and fresh RPMI 2/0 containing MVs at an MOI of 100 MVs per differentiated macrophage was added and incubated for 25 hours. Cells were treated with LPS (1 μg/ml, clone L2654, Millipore Sigma, Burlington, MA) to serve as positive controls. At the end of each treatment period, supernatants were collected, centrifuged for 10 minutes at 4000 rev/min at 4°C and aliquoted. Samples were stored at −80°C until used for downstream analysis.

### Cytokine and Cytotoxicity Analysis

For semiquantitative analysis of cytokines in supernatants from THP-1 cultures, we employed a human cytokine antibody microarray (ab133998, Abcam, Cambridge, UK) according to manufacturer’s instructions as previously described (13). Cells were seeded into 6-well plates at a density of 4 x 10^6^ per well and treated as described above. Membranes were imaged using an Amersham Imager 600 (GE Life Sciences), and densitometry was performed using ImageJ software. Cytokines falling above a fold change of 2 relative to mock treated were considered upregulated and further analyzed. For subsequent analyses of cytokine production, caspase-1 activation, and cell death, cells were seeded into 12-well plates at a density of 2 x 10^6^ cells per well and treated as described above. Cytokines with more than a 2-fold change relative to mock-treated cells were verified using a custom ProcartaPlex bead assay (ThermoFisher, Waltham, MA) as described by the manufacturer. These assays were read and analyzed using a Luminex 200 and Luminex xPONENT v3.1 software, respectively (Luminex Corp., Austin, Texas). Cellular cytotoxicity was assessed using a CyQuant lactate dehytrogenase (LDH) assay (Invitrogen, Waltham, MA) per the manufacturer’s instructions.

### Immunofluorescence Staining and Microscopy Analysis

THP-1 cells were differentiated into 4-well Nunc Lab-Tek II Chamber slides (ThermoFisher, Waltham, MA) at a density of 10^5^ cells per well and differentiated as described above. Cells were treated with either MVs (MOI 100) or LPS (1 μg/mL) for 0.5 or 2 hours and stained for the NF-κB subunit p65 by immunofluorescence as described (36). Briefly, the cells were fixed using 4% paraformaldehyde in PBS for 10 minutes, washed three times with ice cold PBS, and permeabilized for 10 minutes in 0.2% Triton-X in PBS. Cells were washed three more times in PBS and blocked in 10% goat serum/1% BSA/0.3% Tween in PBS for 20 minutes. Rabbit anti-NF-kB antibody (1:1600; clone D14E12; Cell Signaling Technology, Danvers, MA) was added to cells and incubated overnight at 4°C. Cells were washed three times and incubated with Alexa Fluor Goat anti-rabbit 546nm secondary antibody (10 μg/mL; Invitrogen, Waltham, MA) for 1 hour, and washed again in PBS. Coverslips were mounted using Vectashield DAPI (Vector Laboratories, Inc., Burlingame, CA), and representative images were obtained using a Nikon

Eclipse Ti outfitted with a 20x plan fluor objective. Immunofluorescent microscopy was performed in biological triplicate for each timepoint and treatment.

### Caspase-1 Activity, Responses, and Inhibition

After treatment of THP-1 cells, caspase-1 activity was quantified in supernatants using a commercially available assay (Caspase-GLO 1 Assay; Promega, Madison, WI) according to the manufacturer’s instructions. Caspase-1 activity in supernatants was quantified using a GloMax Navigator (Promega, Madison, WI). To assess the impact of caspase-1 on MV-induced IL-1β production, PMA-differentiated THP-1 cells were seeded into 12-well plates and pretreated with 50μM of the caspase-1 inhibitor, Ac-YVAD-CHO (Cayman Chemical Company, Ann Arbor, MI), or 10 μM of the NLRP3 inhibitor, MCC950 (Invitrogen, Waltham, MA), for 30 minutes. Cells were treated with either LPS or GBS as described above, and IL-1β concentrations were measured using a ProcartaPlex simplex assay (ThermoFisher, Waltham, MA).

### qPCR Analysis

PMA-differentiated THP-1 cells, which were seeded in 12-well plates at a density of 2 x 10^6^ cells per well, were left untreated or treated with either GBS bacteria, LPS, or MVs as described above. After 2 or 4 hours, supernatants were aspirated and cells were lysed by adding 1mL Trizol reagent (Invitrogen, Waltham, MA) and gentle scraping. Samples were stored at −20°C until RNA extraction was performed using Phase Lock gel heavy tubes per manufacturer’s instructions (Quanta Bio, Beverly, MA). RNA was quantified using a Nanodrop 8000 spectrophotometer (Thermo Scientific, Waltham, MA) and stored at −20°C until use. Reverse transcription was performed on 0.5 μg of total RNA using Quantitect Reverse Transcription Kit (Qiagen, Hilden, Germany), and 2 μL of the resulting cDNA was amplified by PCR using TaqMan Universal PCR Master Mix (Applied Biosystems, Waltham, MA) with Taqman probes specific for pro-IL-1β (Assay ID: Hs00174097_m1) and GAPDH (Assay ID: Hs99999905_m1). PCR was performed in a QuantStudio 5 real time thermal cycler for 35 cycles (Applied Biosystems, Waltham, MA).

### Data analysis

Data analysis was performed using RStudio. Shapiro-Wilk tests were used to determine whether data followed a normal distribution. Normally distributed data were analyzed for significance using a two-way analysis of variance (ANOVA), followed by a Tukey HSD *post hoc* test. Alternatively, non-parametric data were analyzed using a Kruskal-Wallis test, followed by Dunn’s *posthoc* test to test for differences between groups. Multiple hypothesis testing was corrected using Benjamini-Hochberg or Bonferroni correction when necessary. The analyses used for individual experiments are denoted in the figure legends for clarity.

## RESULTS

### GBS MVs elicit proinflammatory cytokine responses

We first sought to characterize the cytokine response elicited by MVs from a diverse set of GBS strains representing major STs in clinical circulation. Specifically, we characterized the cytokine response to MVs from an ST-1 strain (GB37), two ST-12 strains (GB653 and GB1455), and one ST-17 strain (GB411). Of note, GB37, GB1455, and GB411 were all isolated from infants with invasive infections, whereas GB653 was isolated from an asymptomatically colonized mother. Human cytokine antibody microarrays revealed that MVs from GB411 and GB653 induced cytokine production from THP-1 macrophages. Of the 80 cytokines and chemokines assayed, 7 were upregulated at least 2-fold in comparison to the untreated cells (Supplemental Figure 1). Cytokines upregulated in responses to MVs included the monocyte and neutrophil chemokines, CCL1, CCL2, CXCL1, CCL20, the pyrogen IL-1β, and the proinflammatory cytokine IL-6 (Figure 1, Supplemental Figure 1). Several cytokines were also induced differentially between the two isolates: MVs from GB411 induced CXCL1, CCL1, and IL-1β more strongly than MVs from GB653 (Figure 1). However, the same trend was not observed when comparing cytokines between bacteria-treated THP-1 cells since GB411 and GB653 elicited similar cytokine responses for each of these targets (Figure 1).

**Figure 1:**
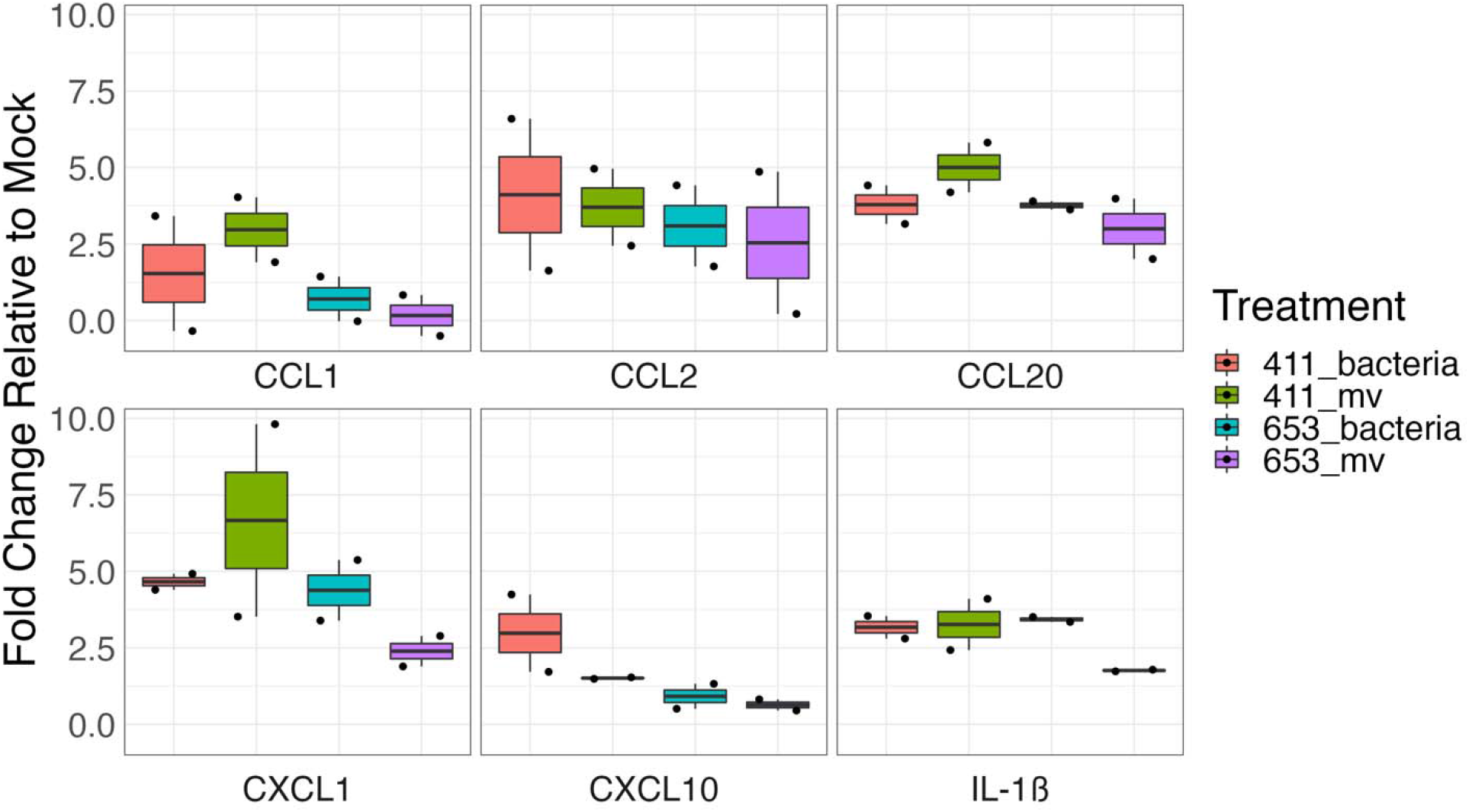
Human antibody cytokine microarray reveals highly upregulated cytokines in response to GBS and GBS MVs. Human cytokine antibody microarrays (Abcam) were probed with supernatants from untreated, bacteria-treated, or MV-treated THP-1-derived macrophages. Membrane densitometry was assessed using ImageJ software. The bacterial strains used are an invasive ST-17 strain (GB411) and a colonizing ST-12 strain (GB653). Shown here are hits of interest that displayed greater than 2-fold change (FC) induction relative to untreated controls in at least one group. Black dots indicate a single biological replicate. n = 2/treatment.

To validate these differences in cytokine production, we used quantitative Luminex-based assays. Consistent with previous results (13), GBS induced a potent proinflammatory response relative to untreated controls (Supplemental Figure 2–3), though IL-6 production remained unchanged by MV exposures (Supplemental Figure 3). Moreover, the MVs induced CCL1, CCL20, CXCL10, CXCL1, and IL-1β, with no differences between the strains from which the MVs were derived (Figure 2). While CCL2 displayed an elevated response relative to mock treatment, this induction was only significant for MVs produced by GB37, GB411, and GB1455.

**Figure 2:**
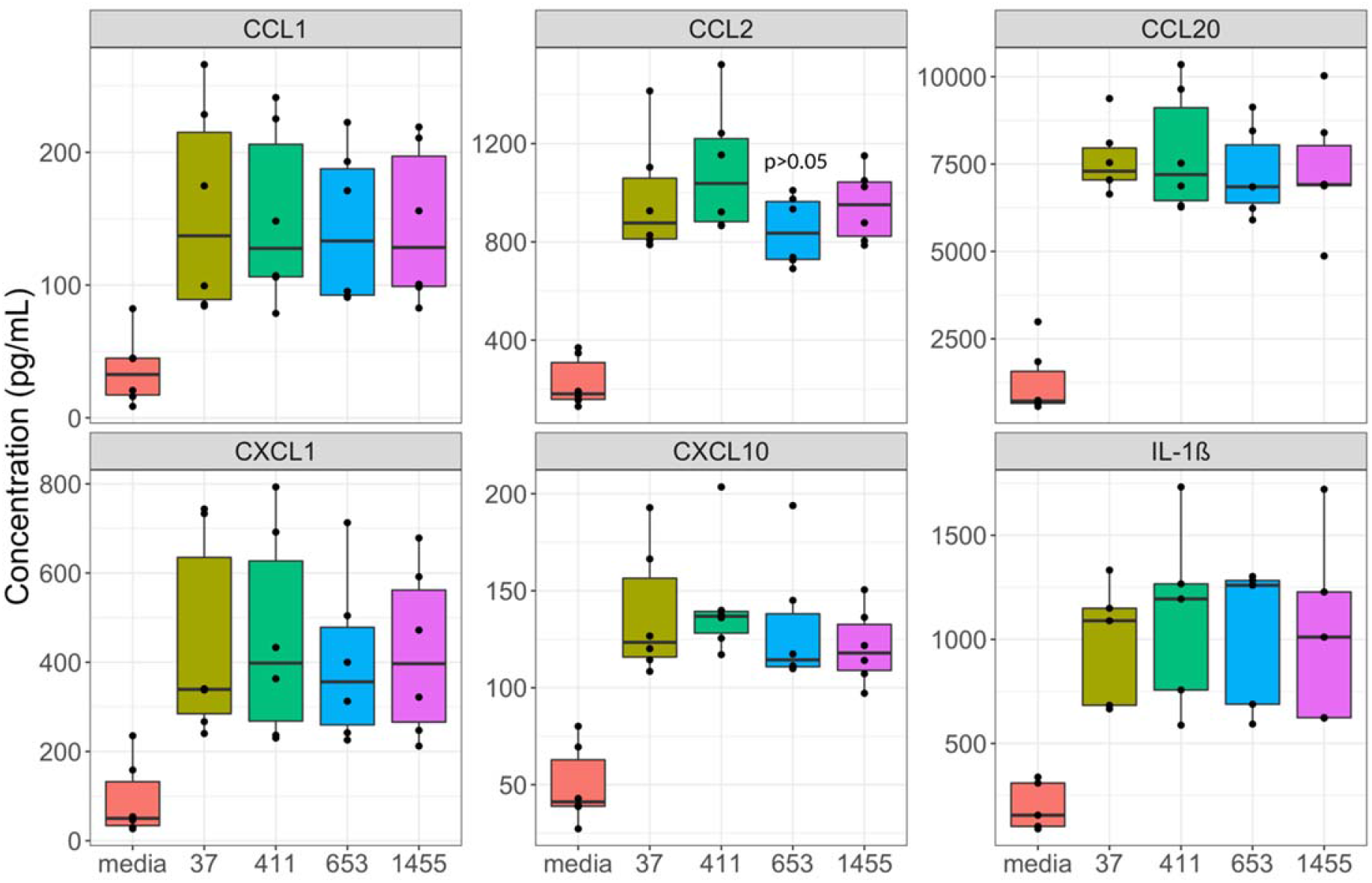
MVs induce proinflammatory cytokine and chemokine responses. Supernatants from THP-1 derived macrophages which were untreated or treated with MVs (MOI 100) for 25 hours were assessed for cytokine production using ProcartaPlex multiplex or singleplex (IL-1B) bead-based assays. Individual black dots indicate a single biological replicate (n = 5-6 for each group). Statistics were determined using either an ANOVA with a Tukey HSD post hoc or a Kruskal Wallis test with a Dunn Test post hoc when appropriate. All comparisons to mock treatment were significant (p<0.05) unless noted with a specific p-value.

Next, we assessed cytotoxicity for all strains examined above using a lactate dehydrogenase activity assay to ensure that these responses were not biased due to differential cell death. In these analyses, we found that bacteria induced moderate cytotoxicity that varied slightly across bacterial strains (Supplemental Figure 4). Notably, GB37 induced significantly more cytotoxicity than GB1455; however, this cytotoxicity was modest. Although low levels of cytotoxicity were observed during MV treatments, with an average of ~6%, the cytotoxicity levels did not vary across MVs produced by the four different GBS strains.

### Membrane vesicles induce caspase-1 activation

Since IL-1β was significantly increased in response to all GBS MVs regardless of the strain, we sought to classify the inflammatory pathways that impact its production. Using the Caspase-GLO 1 assay, we detected caspase-1 activity in our untreated controls as well as our LPS-stimulated control, albeit at a substantially higher magnitude in our LPS control (Supplemental Figure 5). Detectable caspase-1 activity was also observed in response to MVs and the GBS strains, though some differences were noted. Compared to untreated controls, MVs from GB37, GB411, and GB1455 induced the most potent caspase-1 responses, providing confirmation that MVs were capable of inducing caspase-1 activity (Figure 3A, Supplemental Figure 5). Similarly, GB411 bacteria induced a higher degree of caspase-1 activation compared to untreated controls, which is consistent with our previous findings (Supplemental Figure 6).

**Figure 3:**
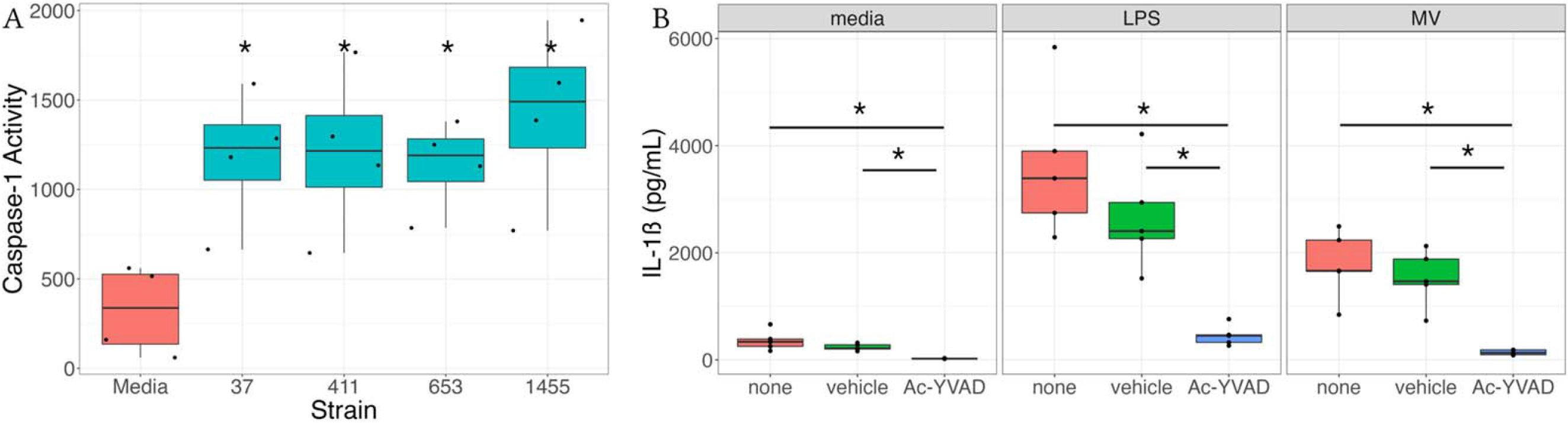
Caspase-1 is critical for the host response to GBS MVs. A.) THP-1 derived macrophages were unstimulated or treated with MVs for 25 hours. Supernatants were then assessed for caspase-1 activity using a caspase-1 GLO assay. Relative light units (RLU) were obtained from a GLO Max Navigator. Data represent the amount of caspase-1 activity (caspase-1 activity = (RLU GLO) – (RLU AC)) from paired samples. B.) THP-1 derived macrophages were pre-treated media, ethanol (vehicle), or Ac-YVAD-CHO for 30 minutes prior to stimulation with LPS, media, or MVs for 25 hours. Supernatants were then assessed for IL-1β concentration using ProcartaPlex bead-based assays. Individual black dots indicate a single biological replicate (n = 4 for each group). Statistical significance is defined as p<0.05 as calculated by ANOVA with a Tukey post-hoc and indicated by (*).

Next, we sought to determine if alternative pathways may be contributing to the conversion of pro-IL-1β to mature active IL-1β. To assess this, we pretreated THP-1 cells with the capsase-1 inhibitor Ac-YVAD-CHO prior to treatment with MVs or LPS for 25 hours. We found that LPS and untreated controls both produced lower amounts of IL-1β when pretreated with Ac-YVAD-CHO compared to the vehicle controls (83% and 90% reduction, respectively); Figure 3B). Furthermore, inhibition of caspase-1 by Ac-YVAD-CHO resulted in almost complete abrogation of MV-stimulated IL-1β secretion (91% reduction) compared to the vehicle control. Importantly, alterations in IL-1β production were not associated with cell death (Supplemental Figure 7). This finding therefore demonstrates that caspase-1 activation is necessary for the maturation of pro-IL-1β to mature IL-1β in response to GBS MVs, regardless of the strain type (Figure 3B).

### NLRP3 is essential for MV mediated IL-1B secretion

Having established that caspase-1 is required for IL-1β maturation, we next investigated the upstream sensor of MVs. Because GBS triggers inflammasome activation via a NLRP3-dependent mechanism, we assessed whether inhibition of NLRP3 could impact caspase-1 activation in response to GBS MVs (37). Notably, inhibition of NLRP3 with the MCC950 inhibitor prevented both MV- and GBS-induced caspase-1 activity (Figure 4A and Supplemental Figure 8). We observed a similar trend in our control cells, demonstrating some baseline inflammasome activity in THP-1 cells; however, the magnitude of inflammasome activation was lower in control groups (Figure 4A). Inhibition of NLRP3 also reduced cytotoxicity for both the GBS bacteria- and MV-treated cells; however, this result was not observed for our untreated controls (Figure 4B).

**Figure 4:**
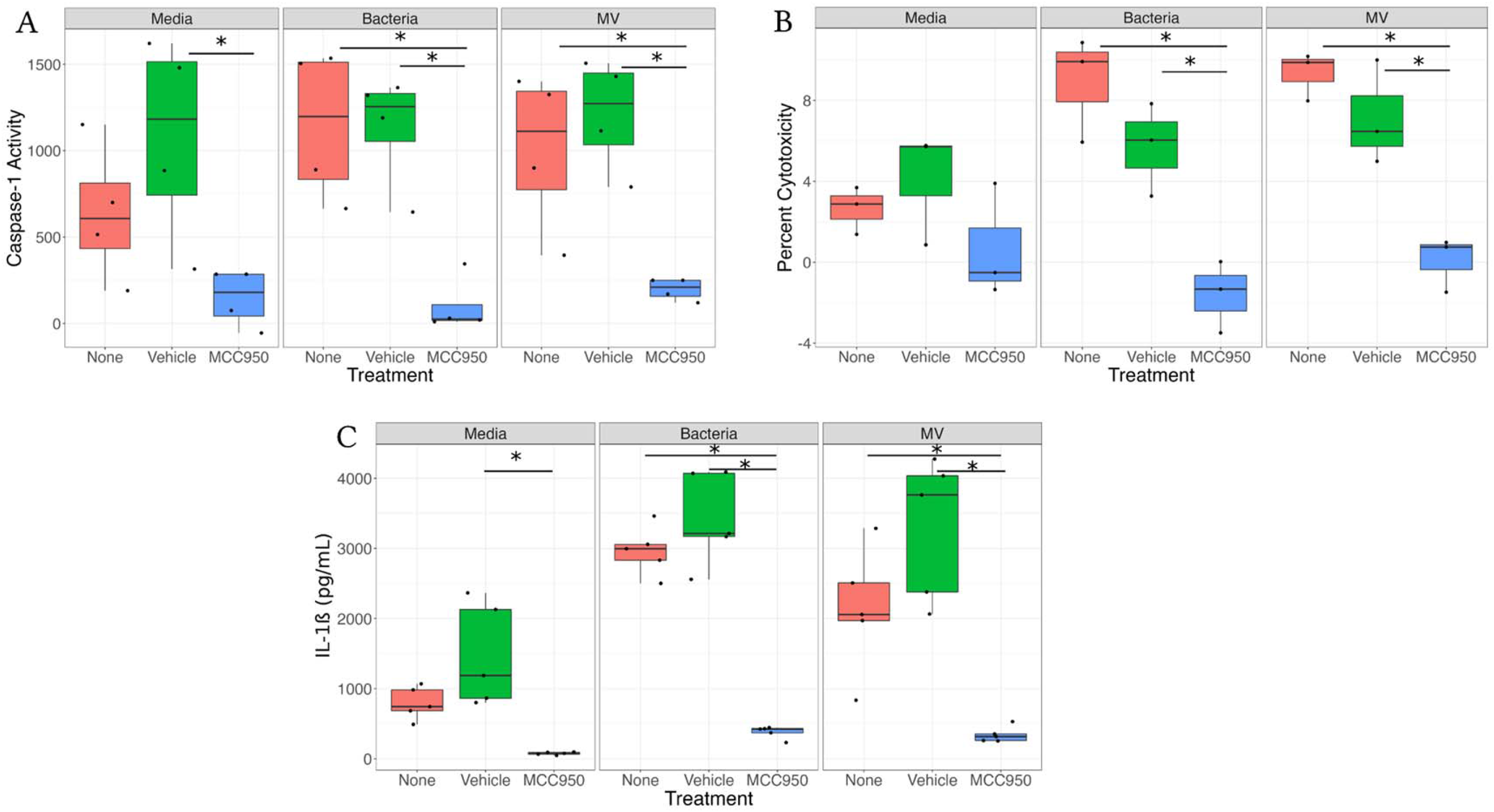
Inhibition of NLRP3 Ablates Caspase-1 Activity in Response to MVs. THP-1s were treated with the NLRP3 inhibitor MCC950 or DMSO for 30 minutes prior to treatment with bacteria, MVs, or media for 25 hours. A.) Caspase-1 activity was determined using the Caspase-1 GLO assay. Caspase 1 activity = ((GLO reagent) – (Ac-YVAD-CHO + GLO Reagent)). Individual points represent individual biological replicates (n = 4 each group). B.) Supernatants were assessed for cytotoxicity using the CyQuant LDH Assay. Individual black dots indicate a single biological replicate (n = 3 for each group). C.) IL-1β contained in supernatants was quantified using ProcartaPlex IL-1β single plex assays. Individual points represent individual biological replicates (n = 5 each group). Statistics were determined using an ANOVA with a Tukey’s HSD post-hoc test. Significance was defined as p<0.05 and denoted with an (*).

Using a similar approach, we also assessed whether NLRP3 impacted secretion of IL-1β from THP-1 cells. In these experiments, we found that inhibition of NLRP3 signaling significantly decreased IL-1β secretion in both the media and LPS controls relative to the vehicle controls (Figure 4C). While the decrease was significant in both groups, the effect was lower for the untreated controls. Moreover, NLRP3 inhibition reduced IL-1β secretion in response to both GBS and the MVs demonstrating that MV-induced IL-1β requires NLRP3 (Figure 4C).

### Membrane vesicles do not trigger transcription activation of pro-IL-1B

We next assessed whether the high levels of IL-1β produced in response to GBS MVs were due to the release of existing pools of pro-IL-1β, or if MVs could directly induce transcription of pro-IL-1β. Using RT-qPCR analysis, we observed no significant increase in pro-IL-1β gene expression relative to untreated cells for LPS, MV, or bacteria treated THP-1 cells at 2 hours post infection (Figure 5A). At 4 hours post infection, however, LPS induced a significant increase in pro-IL-1β gene expression relative to untreated cells, but no similar increases were observed in response to MVs or GBS (Figure 5A). Using immunofluorescence, we similarly found that while LPS rapidly induced the translocation of the NF-κB subunit p65, neither untreated nor MV treated THP-1s induced NF-κB translocation. Notably, MVs induced no NF-κB translocation in response to MVs after a 2-hour exposure (Figure 5B). Similar results were observed at 30 minutes post exposure (Supplemental Figure 9). Together, these data indicate that MVs do not induce a largescale alteration in pro-IL-1β gene expression or NF-κB activation, suggesting that elevated IL-1β secretion is likely due to post-translational regulation.

**Figure 5:**
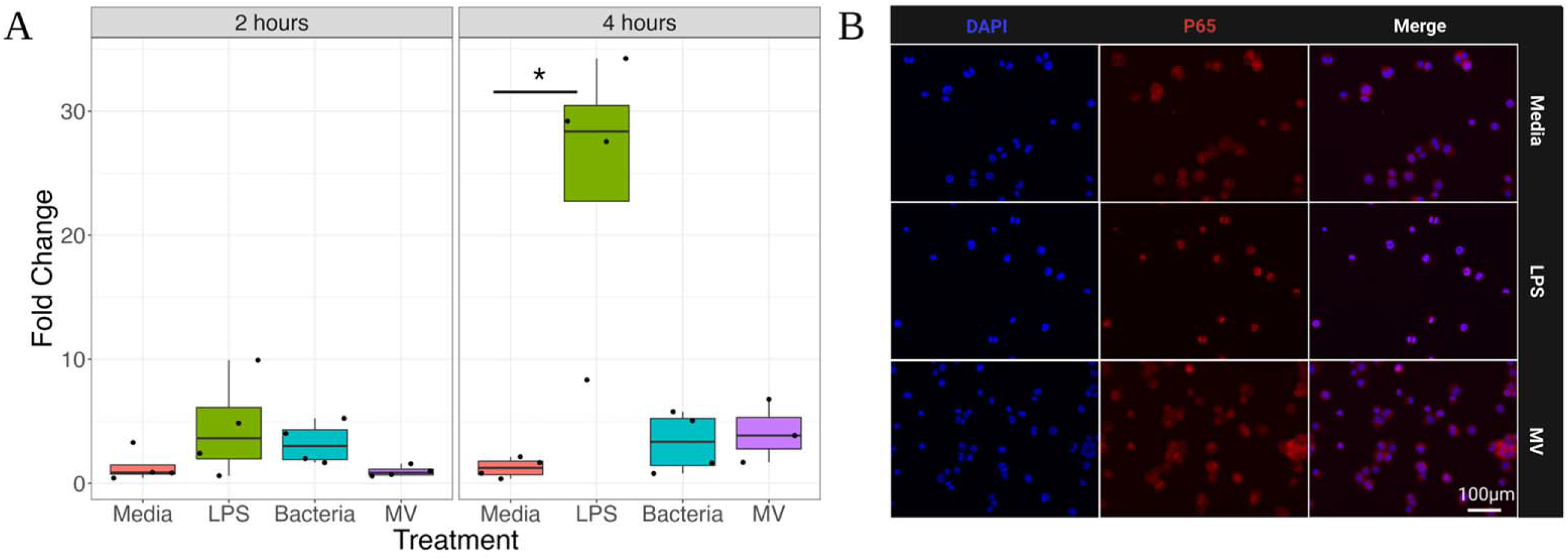
MVs do not prime human macrophages. A.) Expression of pro-IL-1β mRNA from THP-1s treated with bacteria, MVs, LPS, or media was quantified using Taqman probes at 2 and 4 hours after treatment. Fold change values were calculated relative to respective media controls. Each individual dot represents a single biological replicate (n = 4/group). Statistics are calculated using a t-test or Wilcoxson test when appropriate. Significance was defined as p<0.05 and denoted by (*). B.) Differentiated THP-1 derived macrophages were untreated or treated with LPS or MVs for 2 hours prior to fixation and immunofluorescence staining for NF-kB subunit p65 (stained red). Nuclei are stained using DAPI (blue). Shown here are representative images (n = 5) taken at 40x magnification.

## DISCUSSION

Previous studies demonstrated that *in utero* exposure to GBS MVs induced recruitment of neutrophils and lymphocytes into the gestational membranes (17), while MVs induced neutrophil recruitment to the lung in a neonatal sepsis model (19); however, the signals that perpetuate the influx of leukocytes remains unclear. Herein, we have demonstrated that MVs induce expression of proinflammatory cytokines and chemokines in human macrophages *in vitro,* which likely contribute to the inflammatory infiltrate observed *in vivo* (17, 19). Additionally, we found that MVs induce the production of IL-1β by activating pro-IL-1β maturation in an NLRP3 and caspase-1 dependent manner, but independently of NF-κB signaling.

By expanding our current understanding of the cytokine responses towards GBS derived MVs, we have identified the modulators that likely impact the adverse pathologies observed *in vivo.* A previous study demonstrated that the murine chemokine KC, known as CXCL1 in humans, was upregulated in response to GBS MVs (17). In support of these findings, we demonstrate that production of CXCL1 and many additional chemokines are upregulated in human macrophages following challenge with GBS MVs. Notably, CCL1, CCL20, CXCL1, and CXCL10 were all significantly upregulated in response to MVs from four clinical strains. Similarly, the chemokine CCL2 was significantly elevated in response to three of the clinical strains we examined. These chemokines are critical for recruitment of leukocytes to sites of infection, with varying target cell specificities. CXCL1 and CCL20, for example, attract neutrophils (38–40), whereas CCL1 and CCL2 attract monocytes and macrophages (41, 42). Additionally, CCL20 and CXCL10 recruit lymphocytes (40, 43). Unsurprisingly, many of the cytokines have been implicated in GBS-associated disease. For example, CCL20 is upregulated during infection at the blood brain barrier (44). Similarly, CCL2 has been shown to be strongly upregulated during GBS sepsis cases (45). Taken together these data indicate that GBS MVs serve as a critical initiator of disease associated cytokine responses.

Another cytokine that was significantly upregulated in response to MVs was the pyrogen IL-1β, which plays a critical role in the host defense to GBS infections by promoting production of additional neutrophil specific chemokines (20, 46). Although IL-1β does not have direct chemoattractant activity, IL-1β signaling does impact the production of CXCL1 in GBS infections (20). In fact, IL1R knockout mice display reduced neutrophil recruitment and significant increases in mortality when challenged with GBS (46). Given the abundant recruitment of neutrophils and lymphocytes into MV challenged tissues (17), these data provide critical insights into the mechanisms driving this leukocyte infiltration. Although we and others have shown strain variation in IL-1β production in response to whole bacteria (13, 47), here we found that MVs consistently elicited a consistent level of IL-1β from human macrophages, suggesting that it may serve as an important biomarker or possibly a therapeutic target.

Previous studies have highlighted the signaling pathways involved in producing mature IL-1β. Notably, high levels of this cytokine were only produced when both TLR (toll-like receptor) signaling and inflammasome activation occurred (48). TLR signaling occurs when pathogen-associated molecular patterns (PAMPs) engage their cognate receptor (49, 50), which results in the induction of proinflammatory gene expression, including the inactive form of this cytokine, pro-IL-1β (50). Canonically, the induction of pro-IL-1β gene expression depends on translocation of the transcription factor NF-κB, into the nucleus (51).

For pro-IL-1β to be secreted in its mature, active form, a second signal is required. This signal is typically in the form of a danger associated molecular pattern (DAMPs), such as a change in membrane potential due to membrane damage (52, 53). DAMPs are sensed by NLRPs (Nucleotide-binding oligomerization domain, Leucine rich Repeat and Pyrin domain-containing receptors) (52, 54). Once sensed, NLRPs oligomerize with other subunits, forming the inflammasome, (52, 55, 56) which cleaves pro-caspase-1 into its mature, active form (56, 57). Active caspase-1 then cleaves pro-IL-1β, triggering its release (57). This concerted process results in release of stored pools of pro-IL-1β, allowing for rapid immune activation.

Notably, we demonstrate that GBS MVs trigger caspase-1 activation in human macrophages and that the secretion of IL-1β is dependent on caspase-1 activation. Our findings further indicate that MVs do not trigger expression of pro-IL-1β or activation of NF-κB, suggesting that IL-1β production in response to MVs is largely due to post-transcriptional regulatory mechanisms. Interestingly, we also found that caspase-1 activation is ablated in the absence of NLRP3, suggesting that NLRP3 is a sensor of GBS MVs. While previous reports have demonstrated that GBS induces IL-1β production in a NLRP3-dependent manner, this is the first study to demonstrate that GBS MVs contribute to this response (28, 37, 58). Furthermore, we are not aware of any other studies that have identified a pattern recognition receptor capable of sensing GBS MVs. This newfound information may allow for the development of receptor antagonist therapies targeting the NLRP3 dependent recognition of GBS MVs, which could prevent host inflammation and subsequent adverse pregnancy outcomes.

Our data also indicate that inhibition of the NLRP3 inflammasome reduces MV-induced cytotoxicity of macrophages *in vitro.* Several studies have shown that GBS virulence factors, such as hemolysin can induce NLRP3-dependent pyroptosis (37, 58–60). Other studies indicate that GBS mediated pyroptosis is mediated by the activation of the pore forming mediator of pyroptosis, gasdermin D (61, 62). In our examination of THP-1 macrophages, both the MVs and bacteria mediated a modest amount of cell death, which was dependent on the NLRP3 inflammasome, suggesting that MVs can induce pyroptosis, which could suggest gasdermin D activation. While further studies are needed to confirm this hypothesis, the high levels of IL-1β production together with NLRP3 mediated cell death indicate that MVs may be partly responsible for GBS-mediated pyroptosis.

Our analyses also suggest that MVs do not induce pro-IL-1β gene expression. Indeed, while LPS induced a potent upregulation of pro-IL-1β by 4 hours post-exposure, we observed no upregulation of pro-IL-1β in response to MVs or bacteria at either timepoint. Additionally, this lack of induction correlated with the activation of NF-κB signaling, which suggests the following: 1) MVs do not overwhelmingly induce the expression of pro-IL-1β; and 2) the upregulation of IL-1β signaling is likely due to the activation of inflammasome signaling in primed macrophages. Taken together, these data indicate that GBS MVs induce the production of IL-1β in primed macrophages, which is likely a conserved feature of GBS MVs.

A prior study demonstrated that GBS MVs contain active hemolysin and that MV-associated hemolysin exacerbates neonatal sepsis *in vivo* (19). Although it was suggested that GBS-mediated caspase-1 induction requires GBS hemolysin (37), our data indicate that MVs from a non-hemolytic strain of GBS (GB0037) still induce a robust IL-1β response and activate caspase-1 (63). This finding indicates that other factors associated with MVs also induce caspase-1 activation. Indeed, use of proteomics in our prior study found that MVs of different genetic backgrounds contained multiple virulence factors that have been linked to inflammatory responses previously (22). Several factors known to promote immune evasion, such as hyaluronidase, sialidase, and C5a peptidase, were present in GBS MVs at variable levels across diverse phylogenetic backgrounds (22). While these factors can diminish host sensing of GBS, other MV-derived factors likely promote these inflammatory responses. Nonetheless, future studies are required to classify the role that these other factors play in activating these signaling cascades.

Despite advancing our current understanding of the host response elicited towards GBS MVs, it is important to recognize the limitations of our study. Although no strain-specific immune responses towards GBS MVs were observed, we only examined 4 distinct clinical isolates that could have limited our ability to detect differences. Furthermore, our cytokine analysis was limited to those included in the antibody microarrays; hence, it is likely that other responses may also be important. Although our results are consistent with previous reports regarding the host response to GBS MVs (17, 19), it is possible that our system lacks the appropriate complexity to fully model the host response to GBS MVs. Indeed, although THP-1 cells have been shown to largely recapitulate the responses elicited from peripheral blood mononuclear cells, the magnitude of their responses can vary between these two systems (64). Furthermore, the use of cells in monoculture does not capture the complexity of the host responses observed *in vivo.* Therefore, future studies using alternative model systems are warranted.

Overall, data from this study enhance our understanding of how GBS MVs promote both adverse pregnancy and neonatal infection outcomes (Figure 6). It has been established that GBS MVs promote adverse outcomes partly by enhancing neutrophil recruitment (17, 19). In conditions such as chorioamnionitis, we suggest that the sensing of MVs by macrophages may promote proinflammatory immune signaling. Consistent with these findings, we have demonstrated that MVs promote the release of many neutrophil recruiting chemokines as well as the pyrogen IL-1β, which are important for neutrophil recruitment that promote tissue damage via net-osis (20, 40, 65, 66). We also demonstrate that the MV-mediated induction of IL-1β is dependent on caspase-1 activation, which further promotes a proinflammatory environment. Through both direct and indirect tissue damage, MVs likely play a role in weakening gestational membranes, inducing chorioamnionitis, and promoting preterm labor due to enhanced induction of these inflammatory responses (Figure 6). Collectively, these findings expand our understanding of how the immune system respond to these bacterial components that contain important virulence factors capable of initiating an inflammatory response. While the specific PAMPs and DAMPs contained in MVs are not known, this study provides a foundation for future studies aiming to classify the specific factors within MVs that trigger these responses.

**Figure 6:**
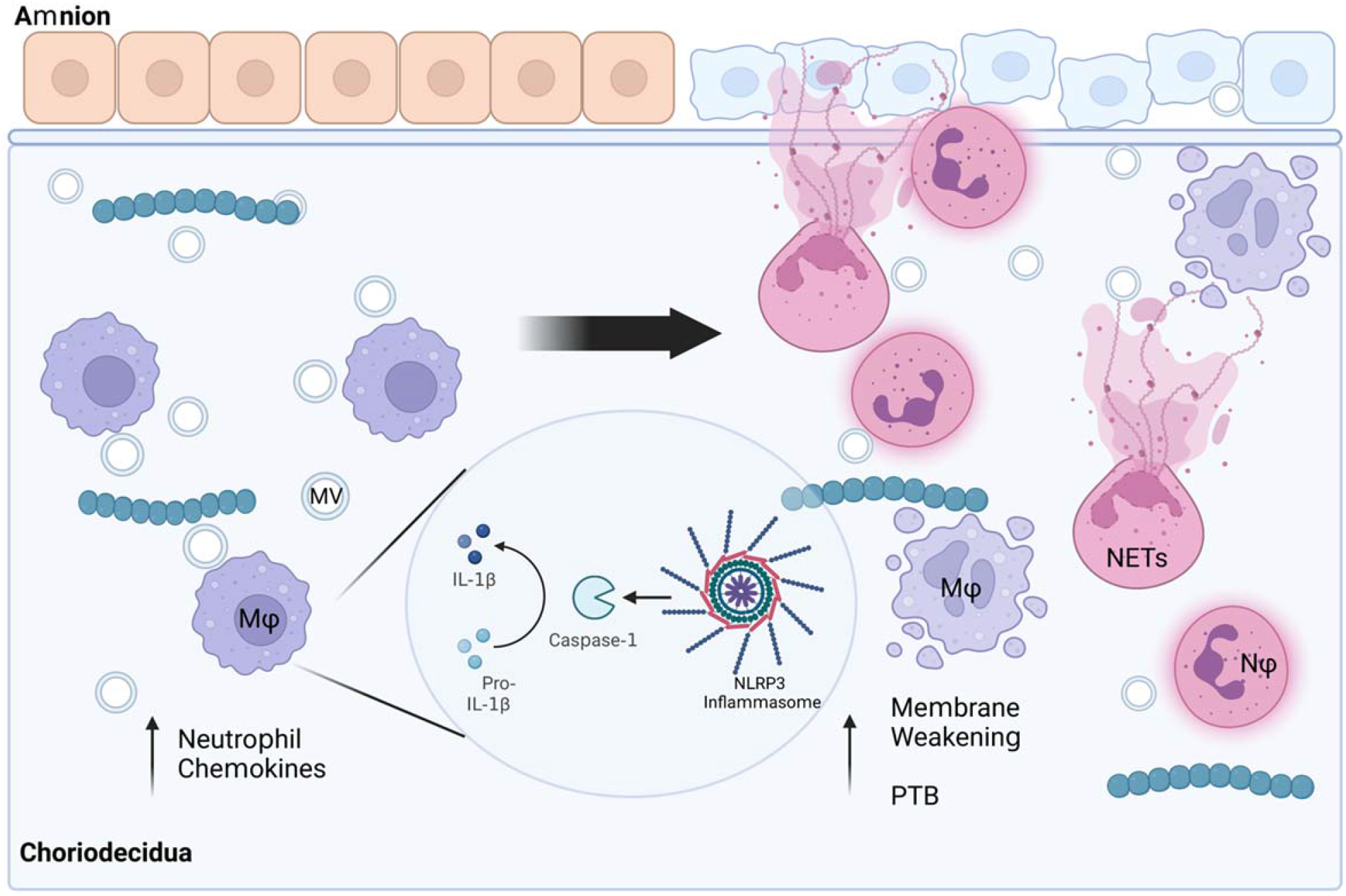
Model of GBS Mediated Chorioamnionitis. GBS is a frequent cause of chorioamnionitis. As sentinel cells at the maternal-fetal interface, macrophages play a critical role in shaping how inflammatory responses are initiated. We show here that macrophages respond to MVs by releasing proinflammatory cytokines and chemokines, many of which recruit neutrophils to the site of infection. Additionally, we show that MVs activate the NLRP3 inflammasome, triggering release of the pyrogen IL-1β. Together these processes promote an influx of neutrophils and leukocytes into the site of infection. In cases such as chorioamnionitis, neutrophils undergo processes including NET-osis, which promote tissue weakening and subsequent preterm birth. Taken together these findings demonstrate mechanistically how MVs may promote preterm birth and chorioamnionitis *in vivo*.

These data illustrate that GBS MVs can induce potent proinflammatory cytokine responses, which is due in part to the activation of the NLRP3 inflammasome. This study advances our understanding of how GBS MVs interact with the host, by identifying the cytokine response towards GBS MVs as well as by identifying NLRP3 as a sensor of MVs. Furthermore, because these cytokine responses are largely conserved across genetically distinct clinical GBS isolates, these responses may represent important targets for immunotherapy or as biomarkers for disease status. Taken together, this study has provided mechanistic insight into the immune response elicited towards GBS MVs.

## Acknowledgments

We would like to thank Dr. H. Dele Davies for sharing the bacterial strains and Drs. Sean L. Nguyen and Soo H. Ahn for helpful conversations and assistance with data analysis. We would also like to thank Dr. Matt Bernard for his assistance with Luminex analyses.

## Author Contributions Statement

CRM, MGP, and SDM designed the study; CRM performed the laboratory work and conducted the analysis; MGP, SDM, DMA, and JGA provided institutional support, guidance and resources, and CRM drafted the manuscript. All authors contributed to and approved of the manuscript content.

---

**Supplemental Figure 1:**
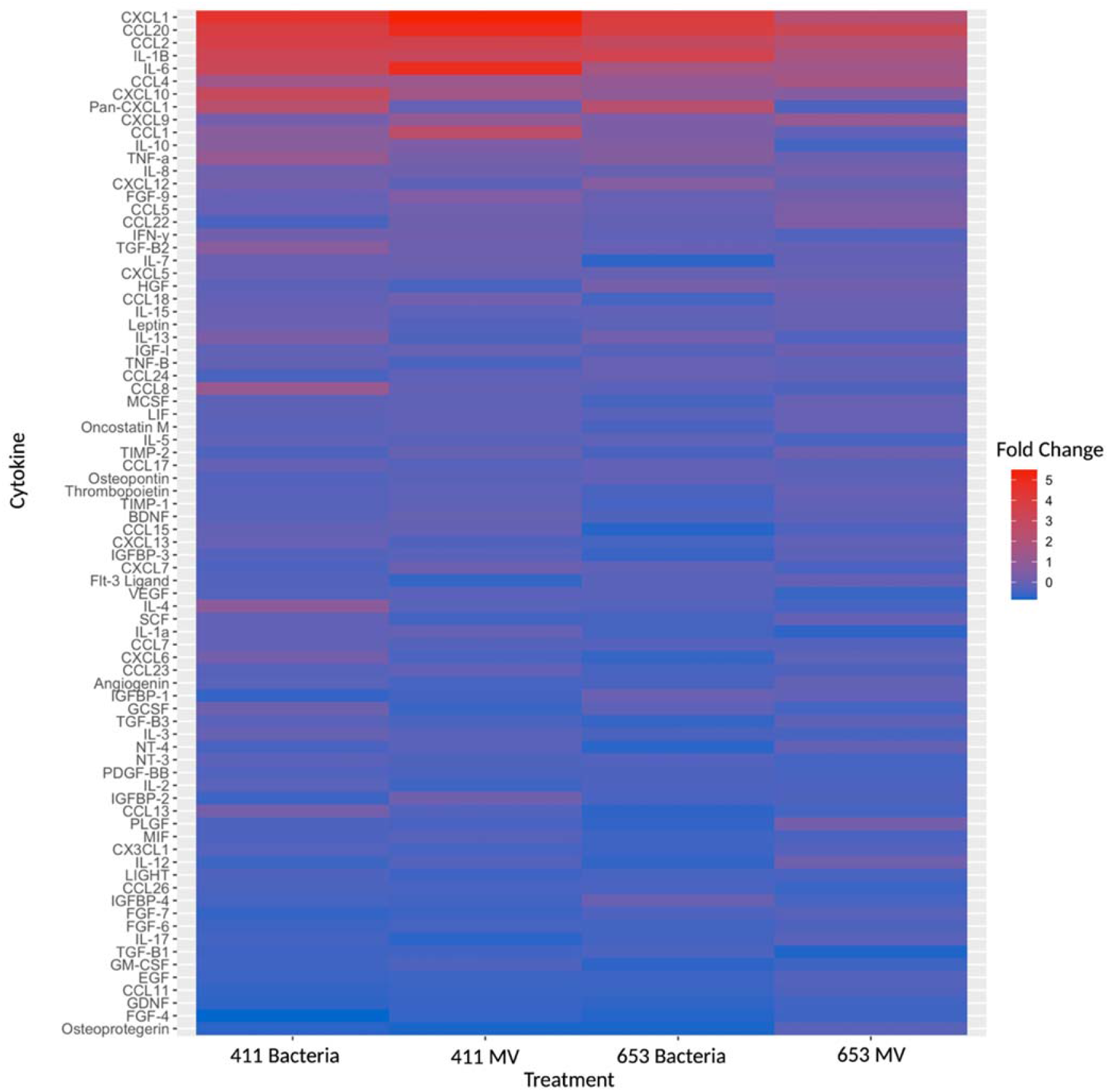
Profiling of cytokine responses elicited towards MVs. THP-1s were treated with bacteria (multiplicity of infection (MOI) = 10) or MVs (MOI 100) for 25 hours prior to supernatant collection. Cytokine production was analyzed using a human cytokine antibody microarray (Abcam). Shown here is semi-quantitative densitometry analysis (ImageJ) of cytokine production for all 80 cytokines examined. Color denotes fold change relative to untreated controls. All groups were performed in biological duplicate. Boxes indicate mean fold change for each condition.

**Supplemental Figure 2:**
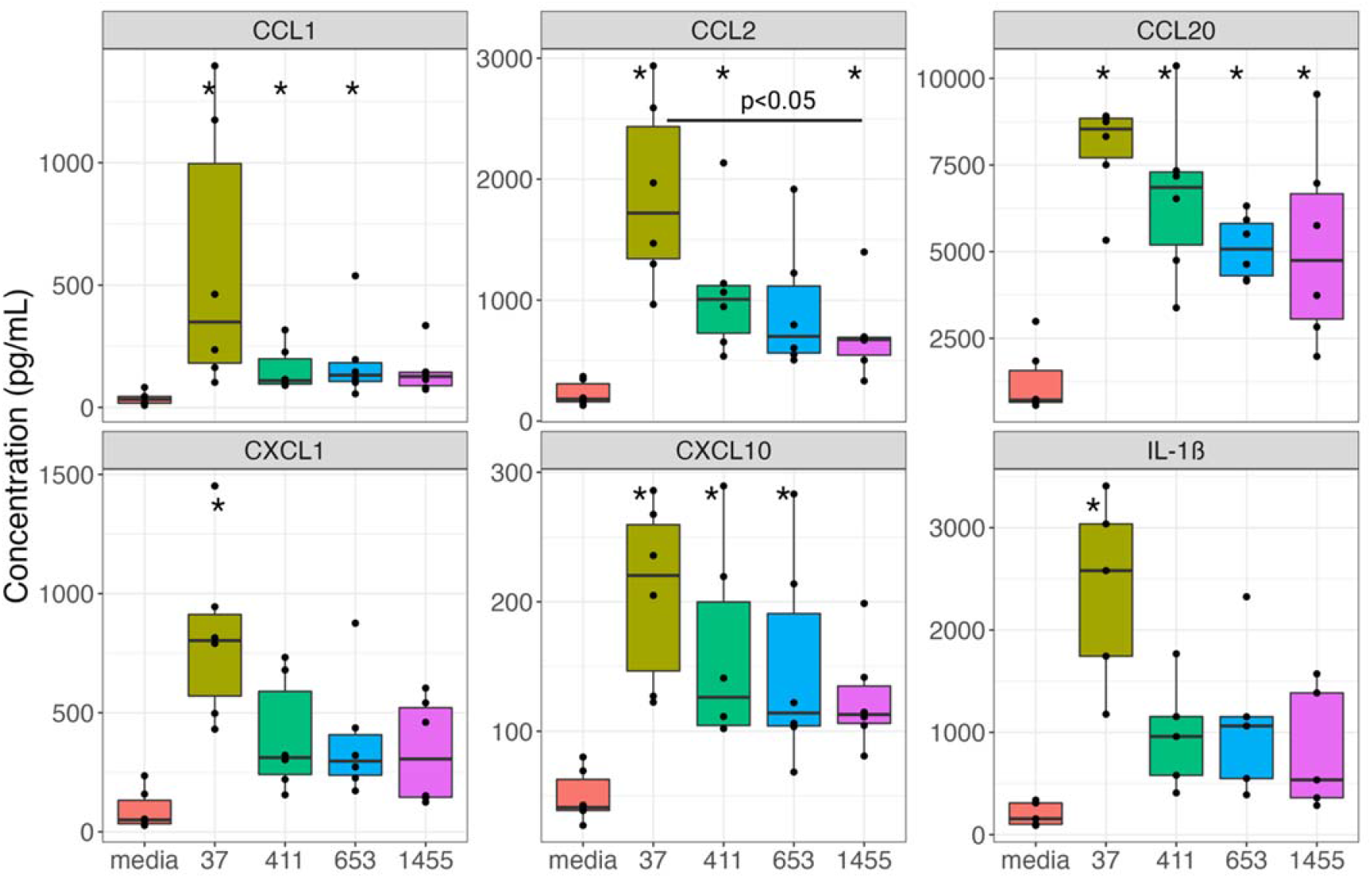
Bacteria elicit proinflammatory immune responses from THP-1 macrophages. Supernatants from THP-1 derived macrophages, which were untreated or treated with bacteria (MOI 10) for 25 hours were assessed for cytokine production using ProcartaPlex multiplex bead-based assays. Each black dot indicates a single biological replicate (n = 5-6 for each group). Data were analyzed by one-way ANOVA with a Tukey HSD post hoc test, or for non-parametric data, a Kruskal Wallis test with a Dunn Test post hoc test. Comparisons with *p* < 0.05 relative to untreated are denoted with (*). Significant differences between strains are denoted with a specific p-value.

**Supplemental Figure 3:**
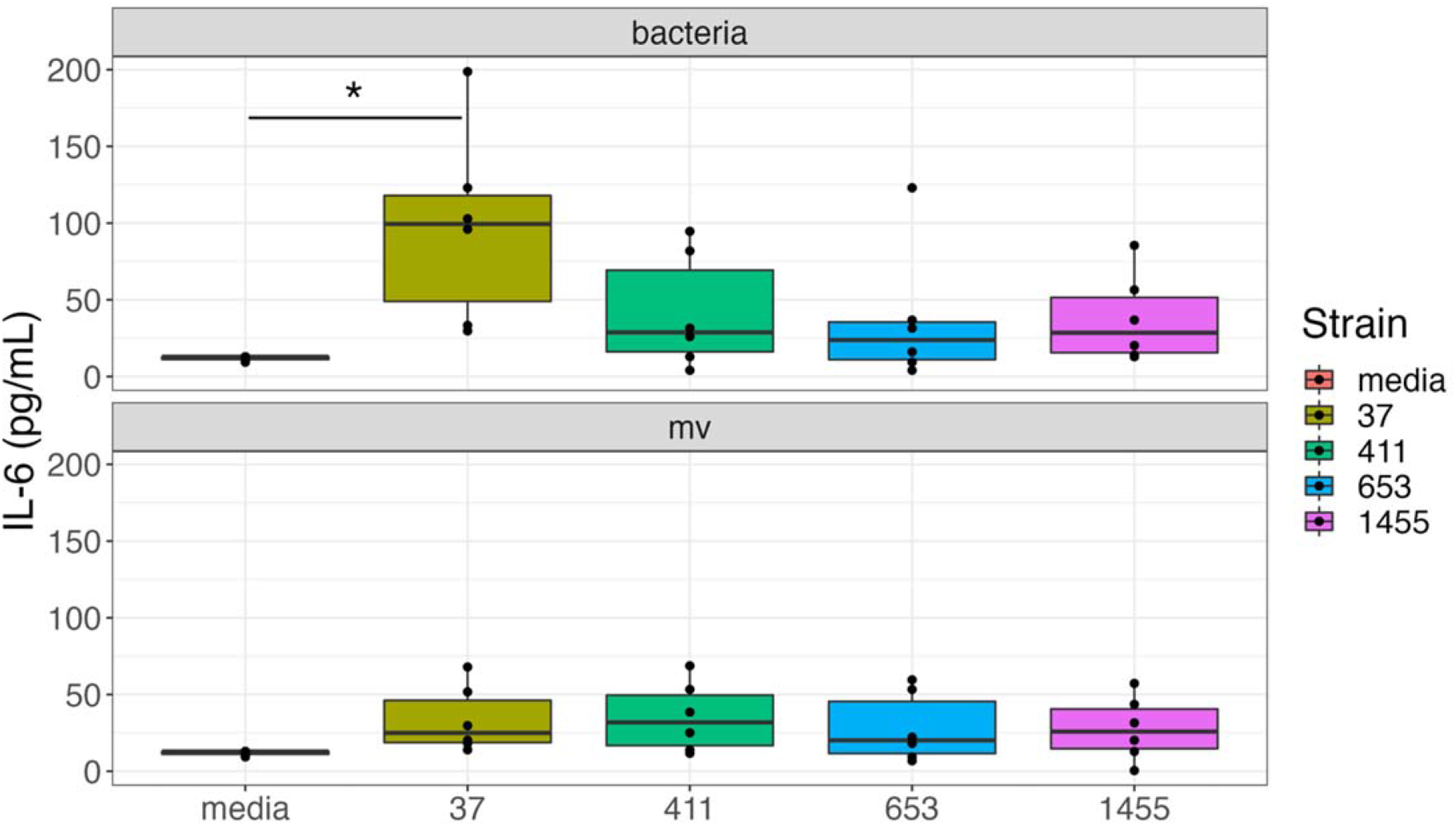
IL-6 is not produced in response to GBS MVs. Supernatants from unstimulated or MV-treated THP-1 derived macrophages were assessed for IL-6 using ProcartaPlex multiplex bead-based assays. Individual black dots indicate a single biological replicate (n = 5-6 for each group). Statistics were determined by one-way ANOVA with a Tukey HSD post hoc, or for non-parametric data, a Kruskal Wallis test with a Dunn post hoc test. Significantly different comparison between groups (P-value < 0.05) are denoted with (*).

**Supplemental Figure 4:**
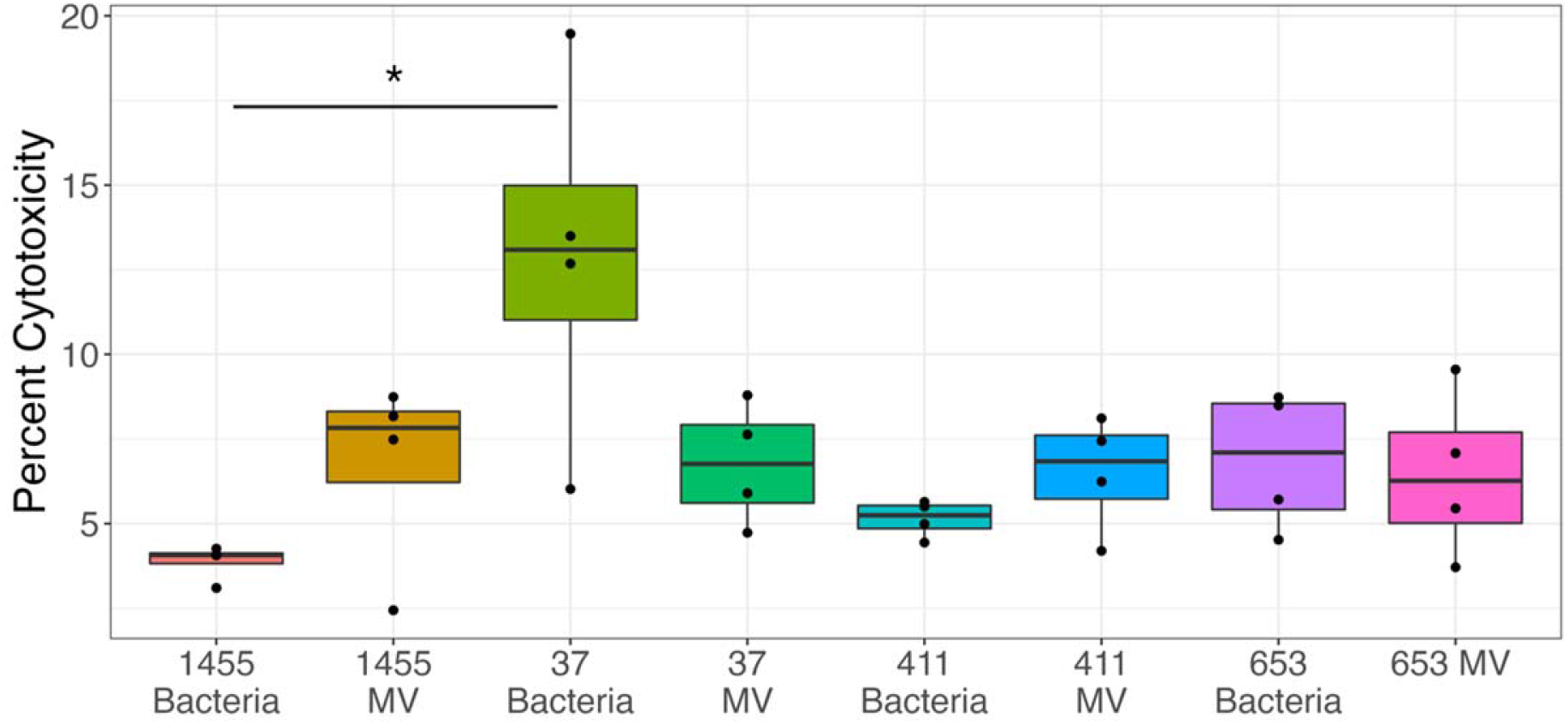
MVs induce a low amount of cell death in THP-1s. THP-1 derived macrophages were unstimulated or treated with MVs for 25 hours. Supernatants were assessed for cytotoxicity using the CyQuant LDH Assay. Percent Cytotoxicity is expressed as a percentage relative to untreated cells. Each black dot represents a single biological replicate (n = 4 /group). Data were anlalyzed using either a one-way ANOVA with a Tukey HSD post hoc test (MV treated groups), or a Kruskal Wallis test with a Dunn Test post hoc (Bacteria-treated groups). Significantly different comparison within groups (P-value < 0.05) are denoted with (*). All other comparisons were not significantly different.

**Supplemental Figure 5:**
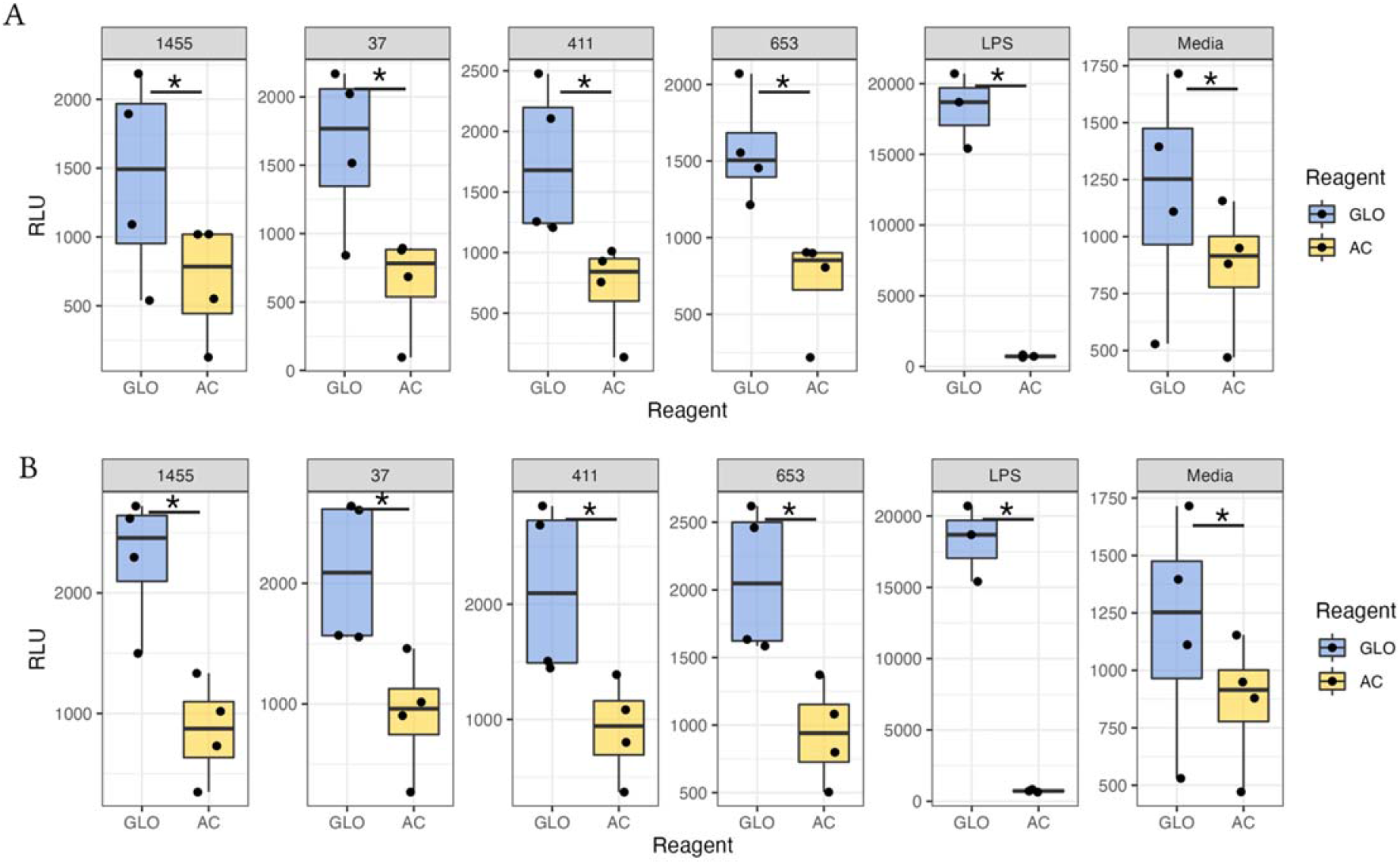
MVs induce caspase-1 activity. Supernatants from THP-1 derived macrophages which were unstimulated or treated with bacteria, MVs for 25 hours. Alternatively, cells were stimulated with LPS for 2 hours. Supernatants were then assessed for caspase-1 activity using a caspase-1 GLO assay. A.) Activity from THP-1s treated with bacteria. B.) Activity from THP-1s treated with MVs. Relative light units (RLU) were determined using a GLO Max Navigator. Individual black dots indicate a single biological replicate (n = 3-4 for each group). Statistics were determined using a two-sided, paired t-test. P-value < 0.05 relative to mock treatment is denoted with a (*).

**Supplemental Figure 6:**
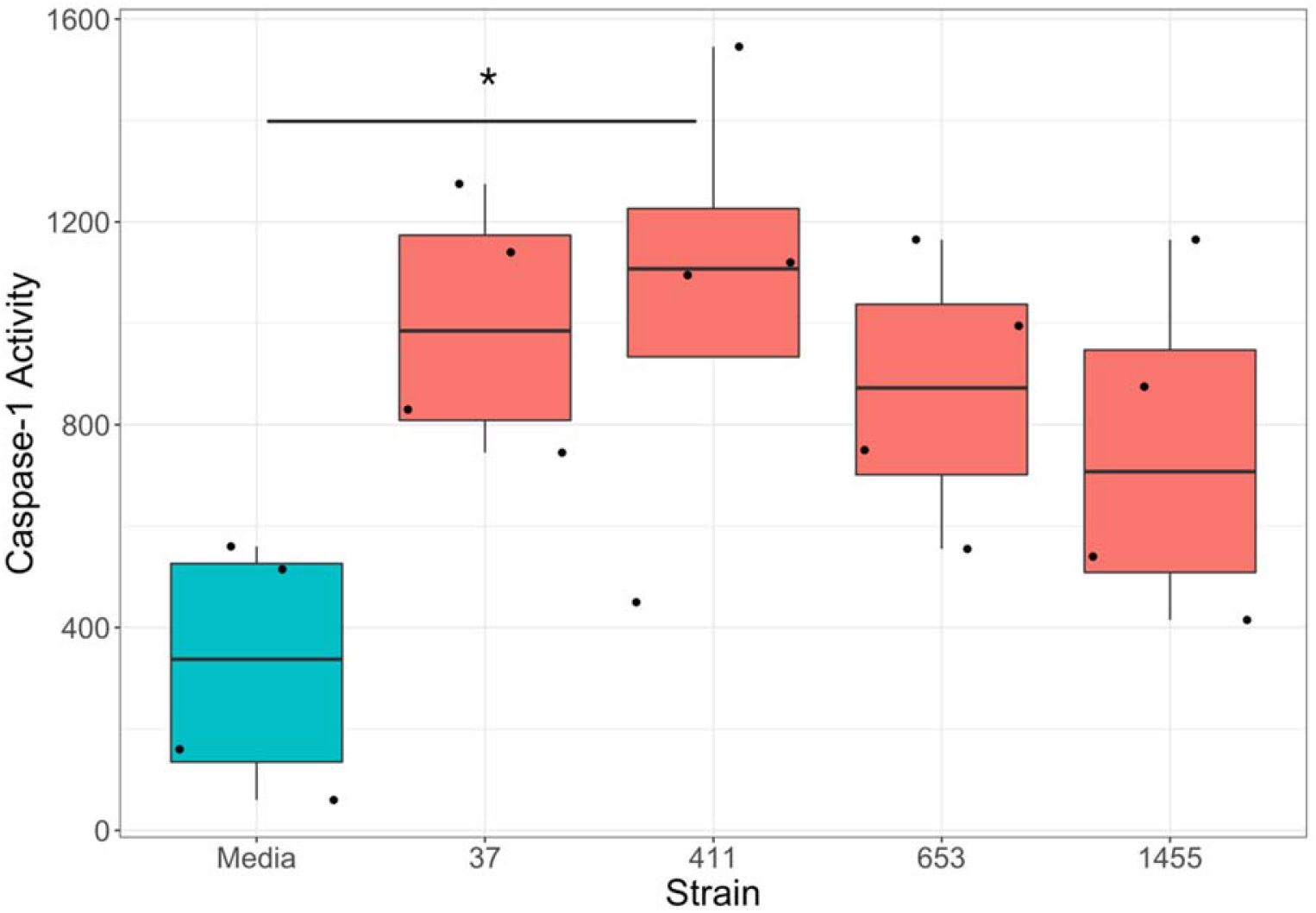
Bacteria induce caspase-1 activation in THP-1s. THP-1 derived macrophages were unstimulated or treated with bacteria for 25 hours. Supernatants were then assessed for caspase-1 activity using a caspase-1 GLO assay. Data represent the amount of caspase-1 activity (caspase-1 activity = (RLU GLO) – (RLU AC)) from paired samples. Individual black dots indicate a single biological replicate (n = 4 for each group). Statistical significance is defined as p<0.05 as calculated by ANOVA with a Tukey post-hoc and indicated by (*).

**Supplemental Figure 7:**
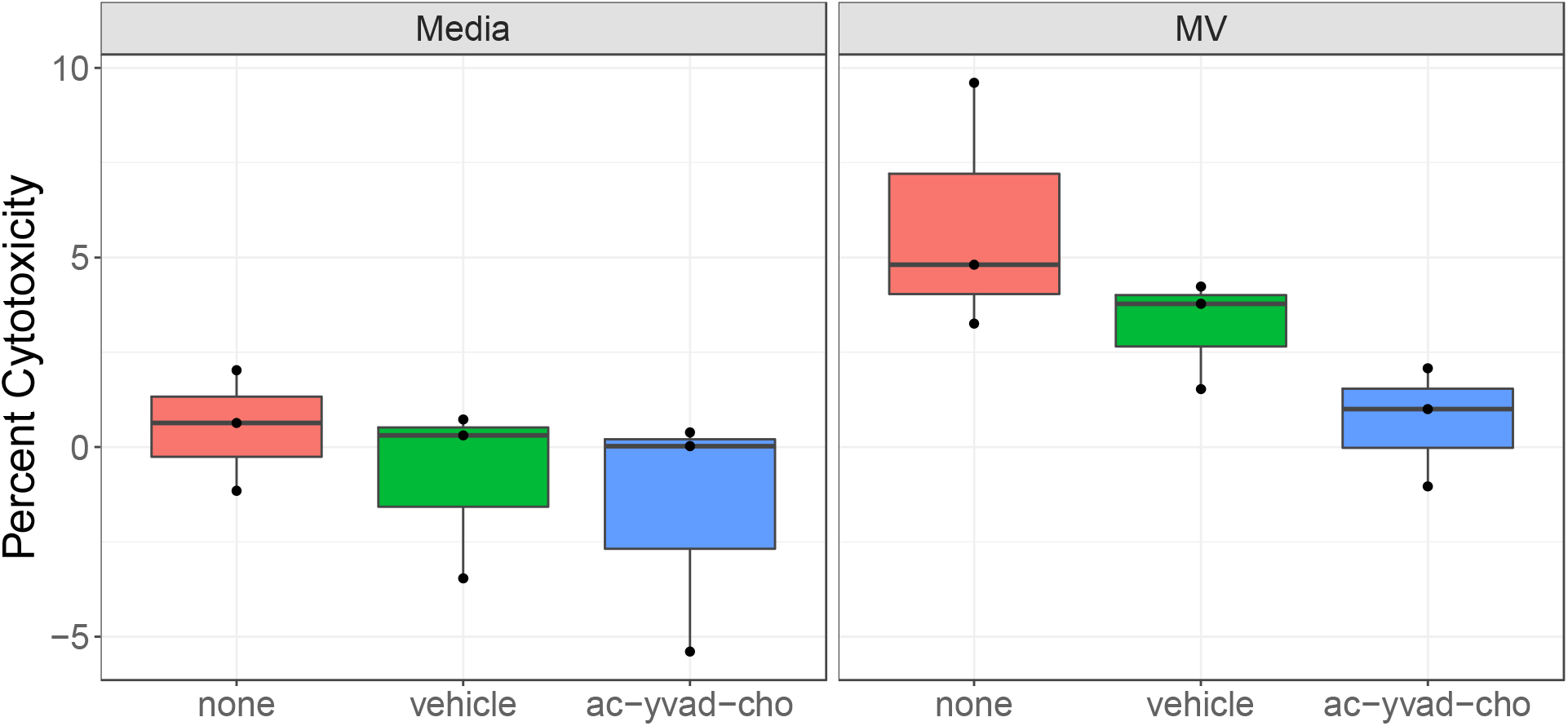
Caspase-1 inhibition does not impact cell death responses. THP-1s were untreated, treated with ethanol, or Ac-YVAD-CHO for 30 minutes. Supernatants from THP-1 derived macrophages, which were subsequently unstimulated or treated with MVs for 25 hours were assessed for cytotoxicity using the CyQuant LDH Assay. Individual black dots indicate a single biological replicate (n = 3 for each group). Statistics were determined using either an ANOVA with a Tukey HSD post hoc. No significant difference relative to non-pretreated cells were detected for either group.

**Supplemental Figure 8:**
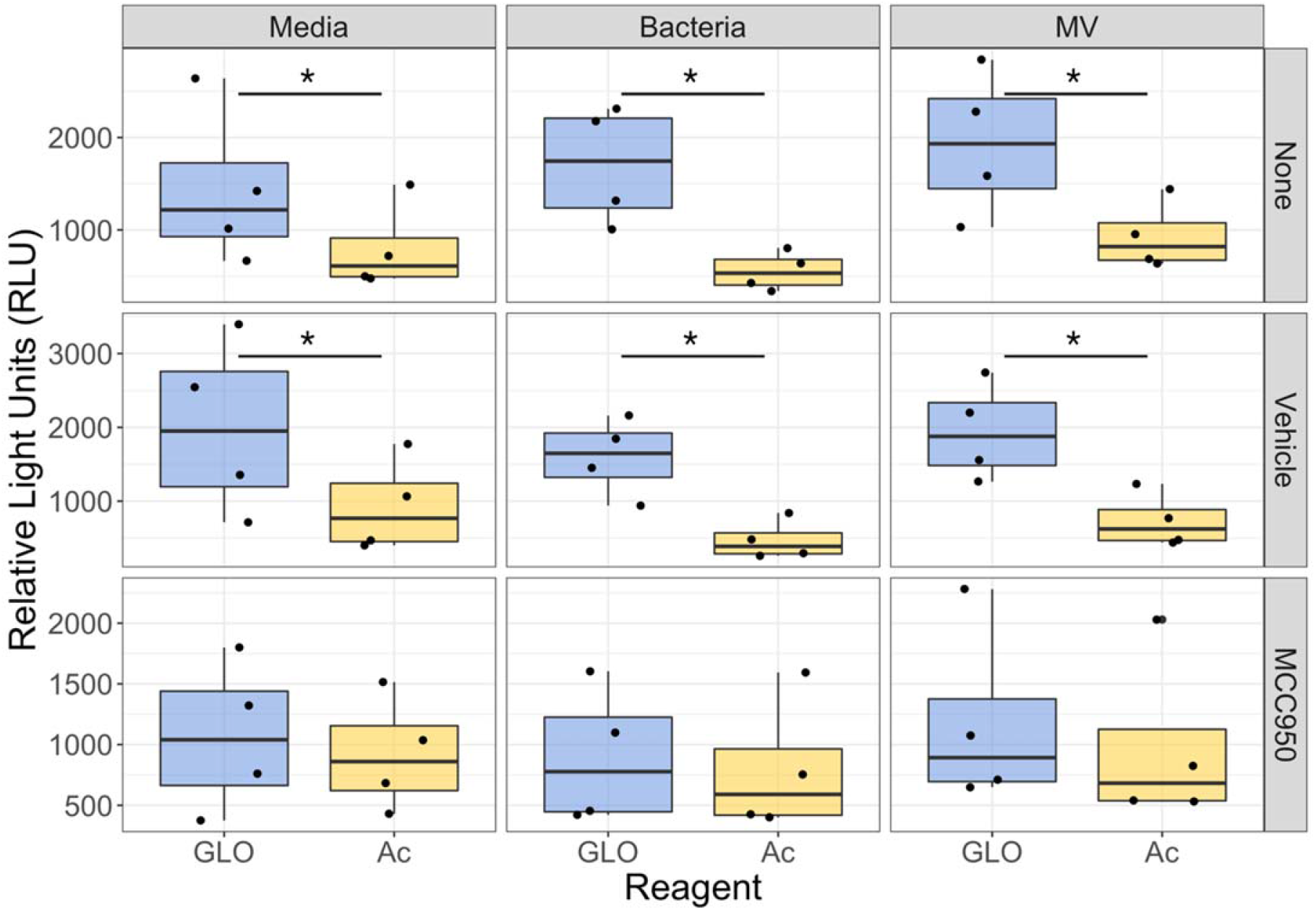
Inhibition of NLRP3 prevents Caspase-1 Activation in Response to MVs. THP-1s were treated with the NLRP3 inhibitor MCC950 prior to treatment with bacteria, MVs, or media. Caspase-1 activity was determined using the Caspase-1 GLO assay. Individual points represent individual biological replicates (n = 4 each group). Statistics were determined using an ANOVA with a Tukey’s HSD post-hoc test. Significance was defined as p>0.05 and denoted with an (*).

**Supplemental Figure 9:**
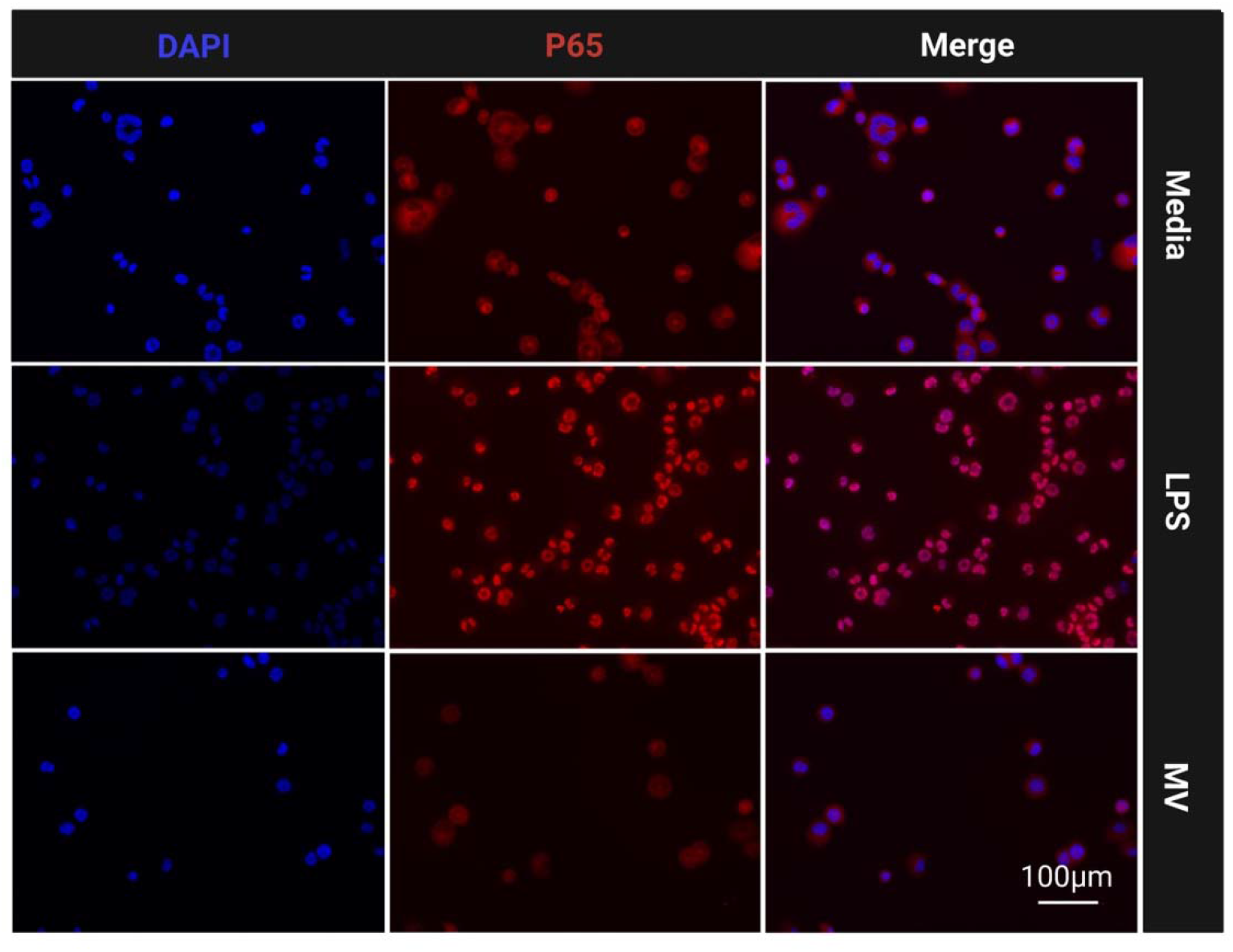
MVs do not prime human macrophages. Differentiated THP-1 derived macrophages were untreated or treated with LPS or MVs for 30 minutes prior to fixation and immunofluorescence staining for NF-kB subunit p65 (stained red). Nuclei are stained using DAPI (blue). Shown here are representative images (n = 5) taken at 40x magnification.

## REFERENCES

1. Verani JR, McGee L, Schrag SJ, Division of Bacterial Diseases NCfIaRD, C.nters for Disease Control and Prevention (CDC). 2010. Prevention of perinatal group B streptococcal disease--revised guidelines from CDC, 2010. MMWR Recomm Rep 59:1–36.

2. Doran KS, Nizet V. 2004. Molecular pathogenesis of neonatal group B streptococcal infection: no longer in its infancy. Mol Microbiol 54:23–31.

3. Bae GE, Yoon N, Choi M, Hwang S, Hwang H, Kim JS. 2016. Acute Placental Villitis as Evidence of Fetal Sepsis: An Autopsy Case Report. Pediatr Dev Pathol 19:165–8.

4. Anderson BL, Simhan HN, Simons KM, Wiesenfeld HC. 2007. Untreated asymptomatic group B streptococcal bacteriuria early in pregnancy and chorioamnionitis at delivery. Am J Obstet Gynecol 196:524.e1–5.

5. Jones N, Bohnsack JF, Takahashi S, Oliver KA, Chan MS, Kunst F, Glaser P, Rusniok C, Crook DW, Harding RM, Bisharat N, Spratt BG. 2003. Multilocus sequence typing system for group B streptococcus. J Clin Microbiol 41:2530–6.

6. Lin FY, Whiting A, Adderson E, Takahashi S, Dunn DM, Weiss R, Azimi PH, Philips JB, Weisman LE, Regan J, Clark P, Rhoads GG, Frasch CE, Troendle J, Moyer P, Bohnsack JF. 2006. Phylogenetic lineages of invasive and colonizing strains of serotype III group B Streptococci from neonates: a multicenter prospective study. J Clin Microbiol 44:1257–61.

7. Luan SL, Granlund M, Sellin M, Lagergård T, Spratt BG, Norgren M. 2005. Multilocus sequence typing of Swedish invasive group B streptococcus isolates indicates a neonatally associated genetic lineage and capsule switching. J Clin Microbiol 43:3727–33.

8. Poyart C, Réglier-Poupet H, Tazi A, Billoët A, Dmytruk N, Bidet P, Bingen E, Raymond J, Trieu-Cuot P. 2008. Invasive group B streptococcal infections in infants, France. Emerg Infect Dis 14:1647–9.

9. Manning SD, Springman AC, Lehotzky E, Lewis MA, Whittam TS, Davies HD. 2009. Multilocus sequence types associated with neonatal group B streptococcal sepsis and meningitis in Canada. J Clin Microbiol 47:1143–8.

10. Flores AR, Galloway-Peña J, Sahasrabhojane P, Saldaña M, Yao H, Su X, Ajami NJ, Holder ME, Petrosino JF, Thompson E, Margarit Y Ros I, Rosini R, Grandi G, Horstmann N, Teatero S, McGeer A, Fittipaldi N, Rappuoli R, Baker CJ, Shelburne SA. 2015. Sequence type 1 group B Streptococcus, an emerging cause of invasive disease in adults, evolves by small genetic changes. Proc Natl Acad Sci U S A 112:6431–6.

11. Manning SD, Lewis MA, Springman AC, Lehotzky E, Whittam TS, Davies HD. 2008. Genotypic diversity and serotype distribution of group B streptococcus isolated from women before and after delivery. Clin Infect Dis 46:1829–37.

12. Korir ML, Laut C, Rogers LM, Plemmons JA, Aronoff DM, Manning SD. 2017. Differing mechanisms of surviving phagosomal stress among group B Streptococcus strains of varying genotypes. Virulence 8:924–937.

13. Flaherty RA, Borges EC, Sutton JA, Aronoff DM, Gaddy JA, Petroff MG, Manning SD. 2019. Genetically distinct Group B Streptococcus strains induce varying macrophage cytokine responses. PLoS One 14:e0222910.

14. Springman AC, Lacher DW, Waymire EA, Wengert SL, Singh P, Zadoks RN, Davies HD, Manning SD. 2014. Pilus distribution among lineages of group b streptococcus: an evolutionary and clinical perspective. BMC Microbiol 14:159.

15. Springman AC, Lacher DW, Wu G, Milton N, Whittam TS, Davies HD, Manning SD. 2009. Selection, recombination, and virulence gene diversity among group B streptococcal genotypes. J Bacteriol 191:5419–27.

16. Brochet M, Couvé E, Zouine M, Vallaeys T, Rusniok C, Lamy MC, Buchrieser C, Trieu-Cuot P, Kunst F, Poyart C, Glaser P. 2006. Genomic diversity and evolution within the species Streptococcus agalactiae. Microbes Infect 8:1227–43.

17. Surve MV, Anil A, Kamath KG, Bhutda S, Sthanam LK, Pradhan A, Srivastava R, Basu B, Dutta S, Sen S, Modi D, Banerjee A. 2016. Membrane Vesicles of Group B Streptococcus Disrupt Feto-Maternal Barrier Leading to Preterm Birth. PLoS Pathog 12:e1005816.

18. De Paepe ME, Friedman RM, Gundogan F, Pinar H, Oyer CE. 2004. The histologic fetoplacental inflammatory response in fatal perinatal group B-streptococcus infection. J Perinatol 24:441–5.

19. Armistead B, Quach P, Snyder JM, Santana-Ufret V, Furuta A, Brokaw A, Rajagopal L. 2021. Hemolytic Membrane Vesicles of Group B Streptococcus Promote Infection. J Infect Dis 223:1488–1496.

20. Biondo C, Mancuso G, Midiri A, Signorino G, Domina M, Lanza Cariccio V, Mohammadi N, Venza M, Venza I, Teti G, Beninati C. 2014. The interleukin-1β/CXCL1/2/neutrophil axis mediates host protection against group B streptococcal infection. Infect Immun 82:4508–17.

21. Lemire P, Roy D, Fittipaldi N, Okura M, Takamatsu D, Bergman E, Segura M. 2014. Implication of TLR-but not of NOD2-signaling pathways in dendritic cell activation by group B Streptococcus serotypes III and V. PLoS One 9:e113940.

22. McCutcheon CR, Pell ME, Gaddy JA, Aronoff DM, Petroff MG, Manning SD. 2021. Production and Composition of Group B Streptococcal Membrane Vesicles Vary Across Diverse Lineages. Front Microbiol 12:770499.

23. Houser BL. 2012. Decidual macrophages and their roles at the maternal-fetal interface. Yale J Biol Med 85:105–18.

24. Care AS, Diener KR, Jasper MJ, Brown HM, Ingman WV, Robertson SA. 2013. Macrophages regulate corpus luteum development during embryo implantation in mice. J Clin Invest 123:3472–87.

25. Rozner AE, Durning M, Kropp J, Wiepz GJ, Golos TG. 2016. Macrophages modulate the growth and differentiation of rhesus monkey embryonic trophoblasts. Am J Reprod Immunol 76:364–375.

26. Doster RS, Sutton JA, Rogers LM, Aronoff DM, Gaddy JA. 2018. Streptococcus agalactiae Induces Placental Macrophages To Release Extracellular Traps Loaded with Tissue Remodeling Enzymes via an Oxidative Burst-Dependent Mechanism. mBio 9.

27. Thomas JR, Appios A, Zhao X, Dutkiewicz R, Donde M, Lee CYC, Naidu P, Lee C, Cerveira J, Liu B, Ginhoux F, Burton G, Hamilton RS, Moffett A, Sharkey A, McGovern N. 2021. Phenotypic and functional characterization of first-trimester human placental macrophages, Hofbauer cells. J Exp Med 218.

28. Sutton JA, Rogers LM, Dixon BREA, Kirk L, Doster R, Algood HM, Gaddy JA, Flaherty R, Manning SD, Aronoff DM. 2019. Protein kinase D mediates inflammatory responses of human placental macrophages to Group B Streptococcus. Am J Reprod Immunol 81:e13075.

29. Tsuchiya S, Kobayashi Y, Goto Y, Okumura H, Nakae S, Konno T, Tada K. 1982. Induction of maturation in cultured human monocytic leukemia cells by a phorbol diester. Cancer Res 42:1530–6.

30. Eastman AJ, Vrana EN, Grimaldo MT, Jones AD, Rogers LM, Alcendor DJ, Aronoff DM. 2021. Cytotrophoblasts suppress macrophage-mediated inflammation through a contact-dependent mechanism. Am J Reprod Immunol 85:e13352.

31. Davies HD, Adair C, McGeer A, Ma D, Robertson S, Mucenski M, Kowalsky L, Tyrell G, Baker CJ. 2001. Antibodies to capsular polysaccharides of group B Streptococcus in pregnant Canadian women: relationship to colonization status and infection in the neonate. J Infect Dis 184:285–91.

32. Spaetgens R, DeBella K, Ma D, Robertson S, Mucenski M, Davies HD. 2002. Perinatal antibiotic usage and changes in colonization and resistance rates of group B streptococcus and other pathogens. Obstet Gynecol 100:525–33.

33. Nguyen SL, Greenberg JW, Wang H, Collaer BW, Wang J, Petroff MG. 2019. Quantifying murine placental extracellular vesicles across gestation and in preterm birth data with tidyNano: A computational framework for analyzing and visualizing nanoparticle data in R. PLoS One 14:e0218270.

34. Nguyen SL, Ahn SH, Greenberg JW, Collaer BW, Agnew DW, Arora R, Petroff MG. 2021. Integrins mediate placental extracellular vesicle trafficking to lung and liver in vivo. Sci Rep 11:4217.

35. Tsuchiya S, Yamabe M, Yamaguchi Y, Kobayashi Y, Konno T, Tada K. 1980. Establishment and characterization of a human acute monocytic leukemia cell line (THP-1). Int J Cancer 26:171–6.

36. Flaherty RA, Aronoff DM, Gaddy JA, Petroff MG, Manning SD. 2021. Distinct Group B. Infect Immun 89.

37. Costa A, Gupta R, Signorino G, Malara A, Cardile F, Biondo C, Midiri A, Galbo R, Trieu-Cuot P, Papasergi S, Teti G, Henneke P, Mancuso G, Golenbock DT, Beninati C. 2012. Activation of the NLRP3 inflammasome by group B streptococci. J Immunol 188:1953–60.

38. Jin L, Batra S, Douda DN, Palaniyar N, Jeyaseelan S. 2014. CXCL1 contributes to host defense in polymicrobial sepsis via modulating T cell and neutrophil functions. J Immunol 193:3549–58.

39. Ritzman AM, Hughes-Hanks JM, Blaho VA, Wax LE, Mitchell WJ, Brown CR. 2010. The chemokine receptor CXCR2 ligand KC (CXCL1) mediates neutrophil recruitment and is critical for development of experimental Lyme arthritis and carditis. Infect Immun 78:4593–600.

40. Hieshima K, Imai T, Opdenakker G, Van Damme J, Kusuda J, Tei H, Sakaki Y, Takatsuki K, Miura R, Yoshie O, Nomiyama H. 1997. Molecular cloning of a novel human CC chemokine liver and activation-regulated chemokine (LARC) expressed in liver. Chemotactic activity for lymphocytes and gene localization on chromosome 2. J Biol Chem 272:5846–53.

41. Qian BZ, Li J, Zhang H, Kitamura T, Zhang J, Campion LR, Kaiser EA, Snyder LA, Pollard JW. 2011. CCL2 recruits inflammatory monocytes to facilitate breast-tumour metastasis. Nature 475:222–5.

42. Cantor J, Haskins K. 2007. Recruitment and activation of macrophages by pathogenic CD4 T cells in type 1 diabetes: evidence for involvement of CCR8 and CCL1. J Immunol 179:5760–7.

43. Liu M, Guo S, Hibbert JM, Jain V, Singh N, Wilson NO, Stiles JK. 2011. CXCL10/IP-10 in infectious diseases pathogenesis and potential therapeutic implications. Cytokine Growth Factor Rev 22:121–30.

44. Kim BJ, Bee OB, McDonagh MA, Stebbins MJ, Palecek SP, Doran KS, Shusta EV. 2017. Modeling Group B. mSphere 2.

45. Okazaki K, Kondo M, Kato M, Nishida A, Takahashi H, Noda M, Kimura H. 2008. Temporal alterations in concentrations of sera cytokines/chemokines in sepsis due to group B streptococcus infection in a neonate. Jpn J Infect Dis 61:382–5.

46. Biondo C, Mancuso G, Midiri A, Signorino G, Domina M, Lanza Cariccio V, Venza M, Venza I, Teti G, Beninati C. 2014. Essential role of interleukin-1 signaling in host defenses against group B streptococcus. mBio 5:e01428–14.

47. Berner R, Csorba J, Brandis M. 2001. Different cytokine expression in cord blood mononuclear cells after stimulation with neonatal sepsis or colonizing strains of Streptococcus agalactiae. Pediatr Res 49:691–7.

48. Kelley N, Jeltema D, Duan Y, He Y. 2019. The NLRP3 Inflammasome: An Overview of Mechanisms of Activation and Regulation. Int J Mol Sci 20.

49. Akira S, Takeda K. 2004. Toll-like receptor signalling. Nat Rev Immunol 4:499–511.

50. Takeuchi O, Akira S. 2002. MyD88 as a bottle neck in Toll/IL-1 signaling. Curr Top Microbiol Immunol 270:155–67.

51. Baeuerle PA, Baltimore D. 1988. Activation of DNA-binding activity in an apparently cytoplasmic precursor of the NF-kappa B transcription factor. Cell 53:211–7.

52. Zheng D, Liwinski T, Elinav E. 2020. Inflammasome activation and regulation: toward a better understanding of complex mechanisms. Cell Discov 6:36.

53. Yu HB, Finlay BB. 2008. The caspase-1 inflammasome: a pilot of innate immune responses. Cell Host Microbe 4:198–208.

54. Mariathasan S, Weiss DS, Newton K, McBride J, O’Rourke K, Roose-Girma M, Lee WP, Weinrauch Y, Monack DM, Dixit VM. 2006. Cryopyrin activates the inflammasome in response to toxins and ATP. Nature 440:228–32.

55. Sharma D, Kanneganti TD. 2016. The cell biology of inflammasomes: Mechanisms of inflammasome activation and regulation. J Cell Biol 213:617–29.

56. Broz P, Dixit VM. 2016. Inflammasomes: mechanism of assembly, regulation and signalling. Nat Rev Immunol 16:407–20.

57. Kostura MJ, Tocci MJ, Limjuco G, Chin J, Cameron P, Hillman AG, Chartrain NA, Schmidt JA. 1989. Identification of a monocyte specific pre-interleukin 1 beta convertase activity. Proc Natl Acad Sci U S A 86:5227–31.

58. Mohammadi N, Midiri A, Mancuso G, Patanè F, Venza M, Venza I, Passantino A, Galbo R, Teti G, Beninati C, Biondo C. 2016. Neutrophils Directly Recognize Group B Streptococci and Contribute to Interleukin-1β Production during Infection. PLoS One 11:e0160249.

59. Whidbey C, Vornhagen J, Gendrin C, Boldenow E, Samson JM, Doering K, Ngo L, Ezekwe EA, Gundlach JH, Elovitz MA, Liggitt D, Duncan JA, Adams Waldorf KM, Rajagopal L. 2015. A streptococcal lipid toxin induces membrane permeabilization and pyroptosis leading to fetal injury. EMBO Mol Med 7:488–505.

60. Gupta R, Ghosh S, Monks B, DeOliveira RB, Tzeng TC, Kalantari P, Nandy A, Bhattacharjee B, Chan J, Ferreira F, Rathinam V, Sharma S, Lien E, Silverman N, Fitzgerald K, Firon A, Trieu-Cuot P, Henneke P, Golenbock DT. 2014. RNA and β-hemolysin of group B Streptococcus induce interleukin-1β (IL-1β) by activating NLRP3 inflammasomes in mouse macrophages. J Biol Chem 289:13701–5.

61. Dubois H, Sorgeloos F, Sarvestani ST, Martens L, Saeys Y, Mackenzie JM, Lamkanfi M, van Loo G, Goodfellow I, Wullaert A. 2019. Nlrp3 inflammasome activation and Gasdermin D-driven pyroptosis are immunopathogenic upon gastrointestinal norovirus infection. PLoS Pathog 15:e1007709.

62. Swanson KV, Deng M, Ting JP. 2019. The NLRP3 inflammasome: molecular activation and regulation to therapeutics. Nat Rev Immunol 19:477–489.

63. Gendrin C, Vornhagen J, Armistead B, Singh P, Whidbey C, Merillat S, Knupp D, Parker R, Rogers LM, Quach P, Iyer LM, Aravind L, Manning SD, Aronoff DM, Rajagopal L. 2018. A Nonhemolytic Group B Streptococcus Strain Exhibits Hypervirulence. J Infect Dis 217:983–987.

64. Chanput W, Mes JJ, Wichers HJ. 2014. THP-1 cell line: an in vitro cell model for immune modulation approach. Int Immunopharmacol 23:37–45.

65. Sawant KV, Poluri KM, Dutta AK, Sepuru KM, Troshkina A, Garofalo RP, Rajarathnam K. 2016. Chemokine CXCL1 mediated neutrophil recruitment: Role of glycosaminoglycan interactions. Sci Rep 6:33123.

66. Kothary V, Doster RS, Rogers LM, Kirk LA, Boyd KL, Romano-Keeler J, Haley KP, Manning SD, Aronoff DM, Gaddy JA. 2017. Group B. Front Cell Infect Microbiol 7:19.

